# Developmental prosopagnosics have normal spatial integration in posterior ventral face-selective regions

**DOI:** 10.1101/2025.07.25.666588

**Authors:** Daniel A. Stehr, Yiyuan Zhang, Anusha Patgiri, Alexis Kidder, Kendrick Kay, Bradley Duchaine

## Abstract

Population receptive field (pRF) mapping is an influential neuroimaging technique used to estimate the region of visual space modulating the response of a neuronal population. While pRF mapping has advanced our understanding of visual cortical organization, evidence linking variation in pRF properties to behavioral performance remains limited. One of the most compelling pRF-to-behavior relationships has emerged from research into developmental prosopagnosia (DP). Individuals with DP have severe deficits in facial identity recognition sometimes linked to diminished holistic processing of faces. This perceptual deficit could be explained at the neural level by abnormally small pRFs in face-selective regions that restrict spatial integration of the face information. This hypothesis is supported by data from a small group of DPs but needs to be rigorously evaluated in a larger sample. Here, we measured pRF properties in 20 individuals with DP and 20 controls using a stimulus designed to robustly activate both low– and high-level visual areas. Consistent with previous studies, DPs exhibited weaker face selectivity in core ventral face-selective areas. Crucially, however, across the visual processing hierarchy – from early visual cortex, to intermediate visual areas, and face-selective areas – DPs and controls exhibited remarkable similarity in pRF properties, including pRF size, visual field coverage, and the distribution of pRF centers. Using a larger sample and the latest methods for mapping and modeling pRFs, these results challenge theories attributing DP to reduced spatial integration in face-selective regions. This underscores the need to explore alternative neural mechanisms of DP and to re-evaluate links between pRF properties and behavior more broadly.

## Introduction

The receptive field (RF) is a fundamental concept in sensory neuroscience. In visual cortex, the RF of a sensory neuron refers to the region of visual space influencing the response of a neuron ***(Hubel and Wiesel, 1965)***. Nearby points in visual space are mapped to nearby points on the cortical surface forming a continuously varying map of the retinal input (i.e. a retinotopic map). Because neurons with similar RFs tend to be grouped together, human neuroimaging techniques have been developed for studying the aggregate RF properties of populations of neurons, termed the population RF or pRF ***(Dumoulin and Wandell, 2008; Harvey et al., 2013; Nasiotis et al., 2017; Winawer et al., 2013)***. Studies of pRFs measured within the volumetric units recorded by functional magnetic resonance imaging (fMRI) have revealed that a visuospatial framework structures much of the cortex – from early visual areas ***(Benson et al., 2018; Deyoe et al., 1996; Engel et al., 1997; Groen et al., 2022; Sereno et al., 1995)*** to high-level visual regions ***(Brewer et al., 2005; Finzi et al., 2021; Grill-Spector et al., 2018; Groen et al., 2022; Kay et al., 2015; Silson et al., 2015)***, and even other sensory modalities ***(Hagler and Sereno, 2006; Silver and Kastner, 2009; Steel et al., 2024; Szinte and Knapen, 2020; van Es et al., 2019)***.

Though a wealth of studies have produced many detailed descriptive accounts of pRF organization across a variety of brain regions, few studies to date have linked individual variation in pRF properties to variation in behavioral performance. One study found a correlation between individual variation in pRF sizes in early visual cortex and perceptual position discrimination thresholds ***(Song et al., 2015a)***. Another study examining ventral face-selective regions found that pRFs mapped with upright face stimuli were larger and higher in the visual field than those mapped with inverted face stimuli ***(Poltoratski et al., 2021***; but see ***Morsi et al., 2024)***, suggesting that large and foveally positioned pRFs are implicated in the common behavioral advantage for recognizing upright versus inverted faces.

The most striking finding linking variations in pRF properties to behavior, however, has emerged from the study of developmental prosopagnosia (DP; also known as *congenital prosopagnosia*). Individuals with DP have great difficulty recognizing facial identity despite having normal low-level vision, normal intelligence, and no history of brain damage ***(Avidan and Behrmann, 2021; Bate and Tree, 2017; Duchaine, 2011; McConachie, 1976; Susilo and Duchaine, 2013)***. Witthoft and colleagues ***(Witthoft et al., 2016)*** conducted retinotopic mapping in seven DPs and 15 controls and found that the DPs had pRFs that were far smaller and closer to the fovea in ventral face-selective regions. For instance, pRFs in a combined ventral face-selective ROI were approximately three times smaller in DPs compared to controls (control mean±SD = 3.4° ± 3.1°, DP mean±SD = 1.0° ± 0.37°). Similar differences were also seen in intermediate regions including hV4 and VO1, whereas no group differences were detected in early visual cortex. Furthermore, mean pRF sizes in the face-selective areas were positively correlated with performance on the Cambridge Face Memory Test ***(Duchaine and Nakayama, 2006)***.

These findings provide an appealing and tractable hypothesis about the abnormal nature of the computations underlying DP. A long history of perceptual research has characterized normal face processing as heavily dependent on the ability to form holistic representations consisting of facial features and their spatial relationships ***(Levine and Calvanio, 1989; Mckone and Yovel, 2009; Rossion, 2009; Tanaka and Farah, 1993; Yin, 1969; Young et al., 1987)***. Although the term *holistic processing* has been conceptualized in a variety of ways ***(Maurer et al., 2002)***, recent theoretical work has grounded it in an understanding of the receptive field characteristics of face-selective regions ***(Avidan and Behrmann, 2021; Grill-Spector et al., 2018; Witthoft et al., 2016)***. When a face is viewed from a typical distance of one meter, an early visual area, such as V1, would not enable holistic face processing, since a representative pRF from V1 only covers an area equivalent in size to the corner of the eye. In contrast, a pRF in a typical observer from a ventral face-selective area, such as one found on the medial fusiform (mFUS) gyrus, covers nearly the entire face, thus enabling the processing of multiple features in parallel as would be required for holistic processing. Therefore, a parsimonious account of the impairment in DP, and one supported by the data from ***Witthoft et al. (2016)***, is that the scope of the spatial integration window in face-selective regions is too small to permit simultaneous analysis of distal features, a phenomenon leading to the inability to get an overview of the face “as a whole in a single glance” ***(Levine and Calvanio, 1989)***. This interpretation dovetails with anecdotal reports from some DPs who have described relying on slow, compensatory strategies that depend on the piecemeal processing of individual face features ***(Duchaine, 2011)***.

The idea that reduced spatial integration underlies the deficit in DP has gained support ***(Avidan and Behrmann, 2021; Jiahui et al., 2018; Kaiser et al., 2019; Linka et al., 2022; Towler et al., 2018)*** and serves as one of the best examples linking pRF organization to high-level perception. However, several methodological limitations of the original study highlight the need for additional investigations into pRF characteristics in DP. First, in the years since the study was conducted, pRF results, especially in high-level regions, have been shown to depend critically on the type of carrier stimulus used during mapping. For instance, the ventral occipito-temporal reading circuitry showed less visual field coverage and higher variance explained for a word stimulus compared to a black-and– white checkerboard stimulus ***(Le et al., 2017)***. Across another set of studies, ventral face-selective regions were found to have lower/upper visual field biases when pRFs were mapped with scene images ***(Silson et al., 2015)*** but no bias when mapped with cartoon imagery ***(Finzi et al., 2021)***. Therefore, pRF mapping stimuli consisting of images of faces, objects, bodies, and scenes have been increasingly used in an effort to better match the preferences of a broad range of category-selective regions ***(Finzi et al., 2021; Kay et al., 2015; Silson et al., 2015)***. However, ***Witthoft et al. (2016)***, used a black-and-white checkerboard pattern as the mapping stimulus. While this decision is unlikely to have affected pRF estimates in EVC, results in higher-level areas are likely to have been impacted. In addition, the sample size used by ***Witthoft et al. (2016)*** is small by current standards (*N* = 7 DPs) and the findings have not been published in a peer-reviewed journal. Large-scale studies and meta-analyses in developmental dyslexia, a condition with many similarities to DP, indicate that neuroimaging studies with small samples are prone to false positives ***(Jednoróg et al., 2015; Ramus et al., 2018)***, echoing the concerns about reproducibility in the wider neuroscience community ***(Boekel et al., 2015; Button et al., 2013; Poldrack et al., 2017; Szucs and Ioannidis, 2017; Turner et al., 2018)***.

To deepen our understanding of the relationship between pRFs in category-selective areas and behavior as well as the role that spatial integration deficits might play in DP, we measured pRF properties in 20 DPs and 20 controls across early, intermediate, and category-selective areas. For mapping pRFs, we used stimuli containing vivid high-level images that effectively stimulate high-level category-selective regions as well as early visual areas. Moreover, we used the compressive spatial summation (CSS) model for estimating pRFs which is a more accurate model for high-level regions where responses to visual stimuli often sum in a subadditive rather than linear manner ***(Kay et al., 2013a***,b). If DP stems from a deficit in spatial integration affecting face perception, we would expect to observe reduced pRF sizes and restricted visual field coverage in face-selective regions of DPs compared to neurotypical controls.

## Results

To investigate spatial integration in developmental prosopagnosia (DP), we measured population receptive fields (pRFs) in a sample of 20 DPs and 20 controls across early, intermediate, and category-selective visual regions. During the pRF mapping experiment, participants fixated a central dot while variously-sized apertures (bars, wedges, and a disk) occupied portions of the visual field revealing colorful, high-level images flickering at a rate of 5 Hz (Supplemental Figure 5). To ensure stable fixation, participants performed a color change-detection task at fixation, and eye movements were recorded for quality control. The compressive spatial summation (CSS) model was used for estimating pRFs ***(Kay et al., 2013a***,b), which produces estimates of pRF center position (expressed in either Cartesian or polar coordinates), pRF size (defined as 2*σ*/ √*n*), pRF gain, and the strength of the static nonlinear exponent (*n*).

Qualitatively, the pRF mapping procedure yielded the expected pattern of retinotopic polar angle and eccentricity maps in all participants – a hemifield representation along the calcarine sulcus flanked by mirror reversals of polar angle, accompanied by foveal representations near the occipital pole that gradually shift to more peripheral eccentricities anteriorly and medially. These maps served as the basis for defining V1, V2, V3, and hV4 with the dorsal and ventral arms of V1 through V3 combined. Delineation of hV4 followed the procedure outlined in ***Winawer and Witthoft (2017)***. Polar angle maps for each individual in the study are displayed in Supplemental Figures 6 and 7 as they serve to demonstrate that participants maintained fixation and pRF mapping worked appropriately.

Participants underwent functional localizer scans to define category selective regions including face-selective regions on the inferior occipital gyrus (occipital face area or OFA but also referred to as IOG-faces, ***Pitcher et al., 2011)***, posterior fusiform gyrus (pFUS, ***Weiner and Grill-spector, 2010)*** and mid-fusiform gyrus (mFUS, ***Weiner and Grill-spector, 2010)***. Additionally, a region on the parahippocampal gyrus selective for scenes (parahippocampal place area or PPA, ***Epstein and Kanwisher, 1998)*** was included to serve as a high-level control region not involved in the perception of faces. To ensure a fair group comparison because individuals with DP often exhibit category selective responses below typically used thresholds, we adopted the “variable-window” method of ROI definition ***(Jiahui et al., 2018; Norman-Haignere et al., 2013)***. In this method, generously-sized group masks for each respective ROI (see Supplementary Figure 11) were intersected with each participant’s individual statistical map and the top 20% most category-selective voxels within the mask were retained for analysis. This method avoids the subjectivity of manually choosing thresholds and ensures that all participants have equally sized ROIs (i.e., identical voxel counts) prior to pRF fittings.

Compared to controls, DPs had lower face-selectivity in all six face-selective ROIs, with face-selectivity defined as the percent signal change to blocks of faces minus the percent signal change to blocks of objects. These differences reached statistical significance in three areas (right and left OFA, and left pFUS; see Supplemental Table 3). Inspection of t-maps thresholded at *t* > 3.3 (uncorrected, *p* < 0.001) overlaid on the anatomical masks revealed that, despite slightly weaker face selectivity, DPs had detectable clusters of activity near the expected anatomical landmarks which is in keeping with previous findings ***(Avidan and Behrmann, 2009; Dinkelacker et al., 2011; Furl et al., 2011; Jiahui et al., 2018)***. Scene-selectivity did not significantly differ between DPs and controls in left and right PPA.

PRF parameters were fit individually to each voxel’s preprocessed time course. Voxels for which the variance explained by the pRF model (coefficient of determination, *R*^2^) was less than 20% were excluded from further analysis. ROIs from any participant with fewer than 10 voxels remaining after *R*^2^ thresholding were omitted from further analysis as the data was deemed insufficient to fully characterize the retinotopic response. The number of surviving ROIs was similar between DPs and controls (Figure 1C) with the exception of left mFUS, which was present in 18/20 controls versus 14/20 DPs.

**Figure 1.**
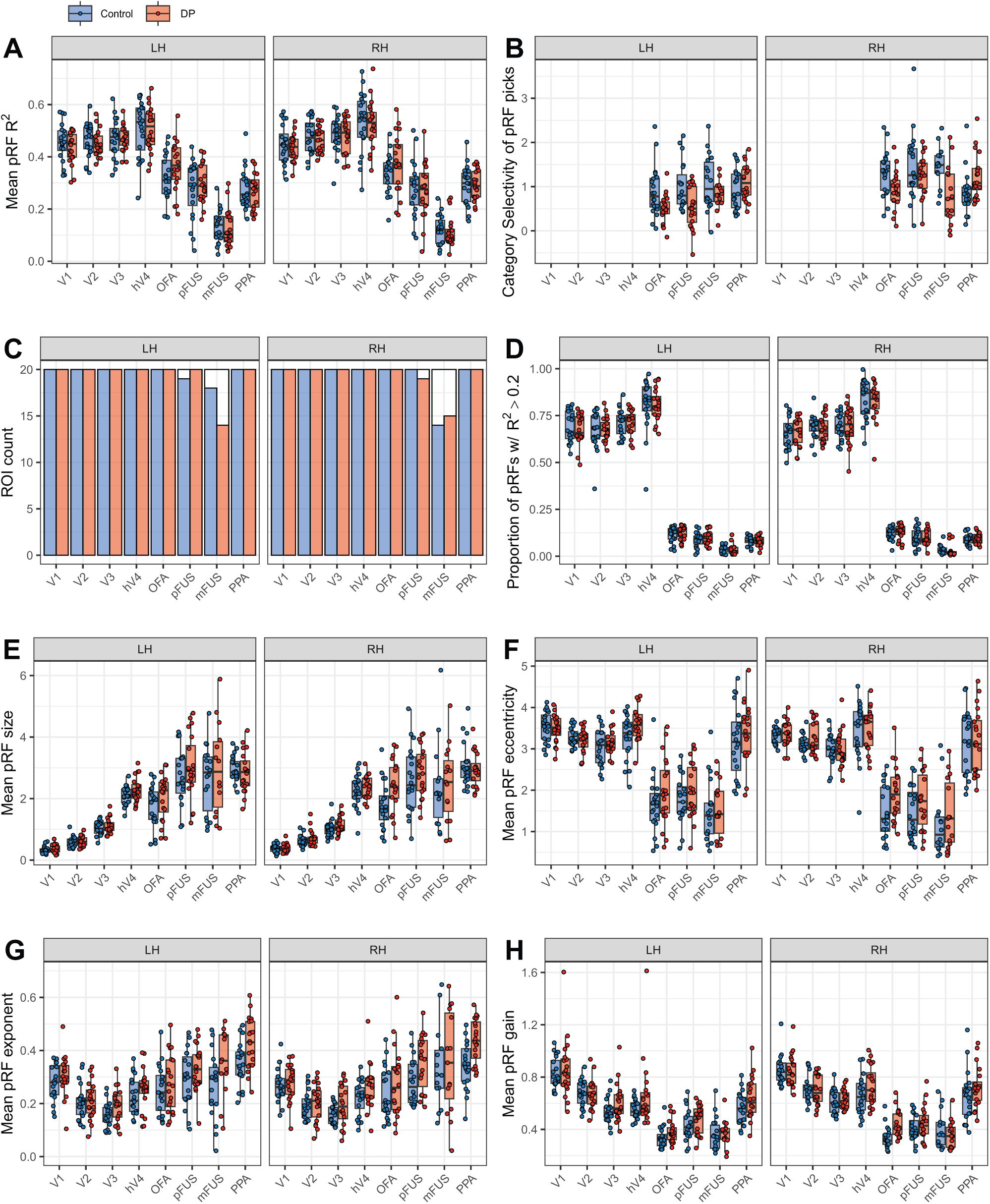
PRF diagnostics and mean parameter values by group (control, DP) and hemisphere (left, right). A) Mean coefficient of determination, *R*^2^, of all pRF fits included in ROIs B) Mean category selectivity of pRF selections from the functional localizer. Category selectivity was defined as the percent signal change for each ROI’s preferred category with respect to percent signal change for objects. Voxels were chosen by selecting the top 20% highest *t* values within anatomical masks. C) Counts of surviving ROIs after pRF fitting by group and hemisphere. An ROI was defined as retinotopically driven, and therefore included for analysis, if more than 10 voxels remained after *R*^2^ thresholding. D) The proportion of voxels that were retained after thresholding on *R*^2^ > .20. E) Mean pRF size, which was defined as *σ*∕ √*n*. F) Mean pRF eccentricity in degrees of visual angle. G) Mean pRF exponent representing the static nonlinearity in the CSS model. H) Mean pRF gain value.

To test for differences between groups (DPs, controls) by hemisphere (right, left) on various measures of pRF properties, a series of linear mixed-effects models (LMMs) were created. LMMs were selected for their ability to: accommodate missing data (e.g. absent ROIs), model the hierarchical/nested structure of the data (such as the fact that right/left hemisphere is nested within each functional ROI and ROIs were nested within subject ***(Etzel et al., 2011)***) and provide robust inference without strict assumptions about sphericity or homogeneity of variances ***(Quené and Van Den Bergh, 2004)***. For each LMM, the random effects component included, at minimum, a random intercept for participant. The fixed effects portion of the model included group, hemisphere, and the interaction between group and hemisphere. In all models, the intercept was mapped to the control group in the left hemisphere. To evaluate the impact of each fixed effect on the dependent variable, likelihood ratio tests were conducted comparing models including the variable (or interaction) in question to a simpler model without it. All statistical analyses were performed in R using the lme4 ***Bates et al. (2015)*** and lmerTest ***Kuznetsova et al. (2017)*** packages.

### Goodness of pRF model fits

We first compared the goodness-of-fit (*R*^2^) of the voxels following pRF fittings. Figure 1A shows that, consistent with previous studies ***(Finzi et al., 2021; Witthoft et al., 2016)***, a higher mean *R*^2^ was observed in the early visual cortex (V1, V2, and V3) and intermediate visual area hV4 than in face-selective areas (OFA, pFUS, mFUS) ***(Finzi et al., 2021; Witthoft et al., 2016)***. This pattern held for both controls (mean ± SE: *EVC* = 53.3±1.29; *hV4* = 55.7±1.75; *ventral face-selective regions* = 41.6±0.81) and DPs (mean ± SE: *EVC* = 51.5±1.01; *hV4* = 53.7±1.72; *ventral face-selective regions* = 42.6±1.24). Similar regional trends were found for the proportion of voxels within an ROI retained after thresholding *R*^2^ at 20% (see Figure 1D). Despite these regional differences in goodness-of-fit, the face-selective ROIs were still strongly retinotopically modulated by the pRF stimulus. Furthermore, across all ROIs examined, DPs and controls did not differ significantly in terms of *R*^2^ nor were there any significant interactions between group and hemisphere (LMMs on *R*^2^ with subject-specific random slopes nested within hemisphere, all *ps* > 0.01, see Supplementary Table 4 for all parameter estimates). The comparable goodness-of-fit values between DPs and controls and strong retinotopic modulation by the pRF stimulus indicate that the data quality was sufficient to support meaningful comparisons of pRF properties in these regions.

### Eyetracking analysis

All participants were instructed on the importance of maintaining steady fixation throughout the entirety of the retinotopic mapping runs. Before beginning the experiment, participants were given ample time to practice fixating while inside the scanner, and eye tracking was performed whenever possible. The experimenters monitored real-time plots of eye positions during data collection and provided feedback in between runs, if needed. One control was eliminated from the study due to drowsiness and inability to maintain adequate central fixation. Another control had one of five retinotopic mapping runs dropped from analysis because the spread of eye positions in that run was unusually large. In total, eye tracking data was successfully recorded for all or some runs of the experiment in 14/20 controls and 14/20 DPs. An LMM on the area of the 95% probability contour of the distribution of eye positions (a summary of bivariate scatter) revealed no significant effect of group (*χ*^2^(1) = 0.40, *p* = 0.529), though there was a main effect of run (*χ*^2^(4) = 16.30, *p* = 0.003), with later runs having more dispersed eye positons. Despite this main effect of run, 95% probability contours were small across the majority of runs revealing that participants, regardless of group, were able to maintain central fixation throughout the retinotopic mapping runs. Heatmaps of eye position recordings for individual participants and runs are displayed in Supplemental Figures 8 and 9 and a group summary is displayed in Supplemental Figure 10.

### DPs do not have restricted visual field coverage in face-selective or intermediate visual regions

Spatial representations within functional ROIs are built up from the responses of many pRFs that vary in their position and size within the visual field. To evaluate how pRFs collectively tile the visual field in DPs and controls, we measured the visual field coverage (VFC) within each ROI. To do this, pRFs were conceptualized as circles of radius 2*σ*/ √*n*, and VFC was computed as the proportion of these circles covering each point in visual space (Figure 2A).

**Figure 2.**
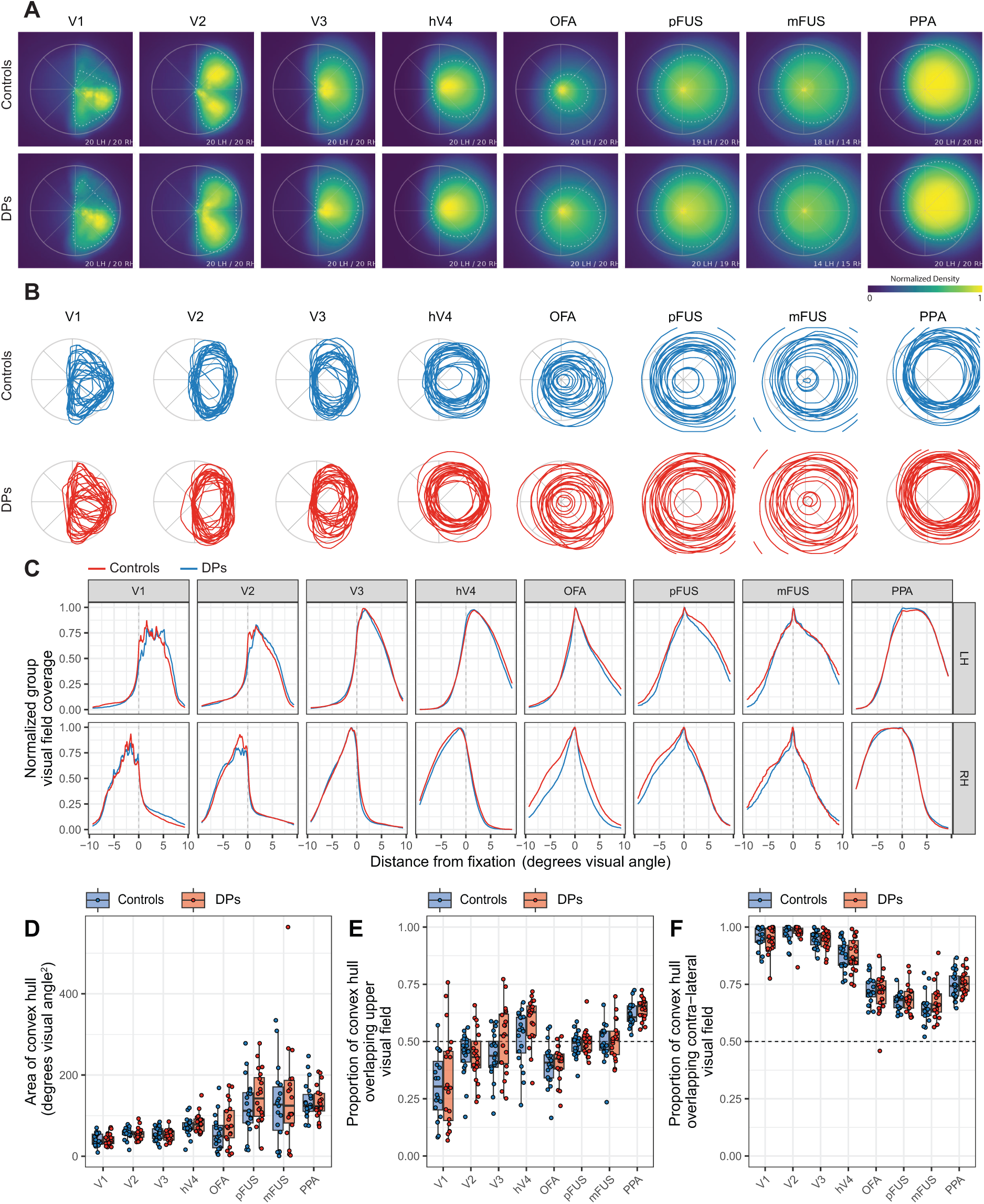
Measures of visual field coverage (VFC). VFC was computed as the proportion of pRFs covering each point in the visual field. A) Group averaged plots of VFC by ROI. Data from the right hemisphere has been reflected across the vertical meridian and combined with data from the left hemisphere. The dotted lines show the convex hull polygon at the 50% density threshold. B) Convex hull polygons, computed at the 50% density threshold, displayed for each individual subject by group and ROI. C) Cuts through the horizontal meridian showing normalized group coverage to the left and right of fixation (at 0 degrees) by hemisphere. D) Total area (in degrees of visual angle squared) for the convex hull polygons in panel B. Datapoints represent individual subjects. E) Proportion of the total convex hull area falling above the horizontal meridian. Values above 0.5 indicate an upper visual field bias and values below 0.5 indicate a lower visual field bias. F) Proportion of the convex hull area falling into the contralateral visual field.

To better quantify variation in VFC across individuals and groups, we computed the convex hull area of the VFC density, which is the area (in degrees of visual angle squared) of the smallest convex polygon wrapping around all points at a given density threshold. For the density threshold, we chose 50% of the normalized density as a robust middle ground. The convex hull area was measured separately for each participant and ROI. Figure 2B displays the convex hull polygons from each participant. In addition, a group-averaged convex hull polygon is superimposed on the visual field coverage density plots in Figure 2A as a dotted line.

In general, the VFC increased as one ascends from V1 through hV4 to the ventral face-selective regions, in accordance with previous studies ***(Benson et al., 2018; Finzi et al., 2021; Kay et al., 2015)***. An LMM with convex hull area as dependent measure and subject-specific random intercepts and fixed effects of ROI (V1, V2, V3, hV4, OFA, pFUS, mFUS, and PPA), group (DP, control), and hemisphere (right, left) revealed a strong main effect of ROI (*χ*^2^(6) = 160.71, *p* < 0.001), indicating that convex hull area was larger for later visual areas. However, no other main effects or interactions were significant (all *ps* > .05).

For all participants, we also computed the proportion of the total convex hull area that fell above the horizontal meridian (upper visual field bias) and the proportion that fell to the contralateral side of the vertical meridian (contralateral visual field bias). An LMM on the proportion of convex hull area above the horizontal meridian revealed a strong main effect of ROI (*χ*^2^(7) = 216.31, *p* < 0.001) but no other significant main effects (including group or hemisphere) and no significant interactions. The regions of V1, V2, and OFA exhibited a lower visual field bias (V1: *t*(39) = –6.56, *p* < 0.001; V2: *t*(39) = –3.53, *p* = 0.001; OFA: *t*(39) = –7.18, *p* < 0.001), and hV4 and PPA exhibited an upper visual field bias (hV4: *t*(39) = 2.94, *p* = 0.006; PPA: *t*(39) = 16.80, *p* < 0.001). Neither pFUS nor mFUS exhibited either an upper or lower visual field bias, which is consistent with several previous reports ***(Finzi et al., 2021; Kay et al., 2015***, but see ***Silson et al., 2015)***. An LMM on the proportion of the convex hull area covering the contralateral visual field revealed a strong significant effect of ROI (*χ*^2^(7) = 132.72, *p* < 0.001) but no other significant main effects or interactions.

Together, these results indicate that, across the visual processing hierarchy, DPs and controls have highly similar visual field characteristics and, notably, DPs do not have reduced coverage of the visual field in any of the ROIs studied, including face-selective ones.

### DPs have normal scaling between eccentricity and pRF size

Previous studies have found a strong linear relationship between pRF eccentricity and pRF size that spans early, intermediate, and ventral (category-selective) visual areas ***(Finzi et al., 2021; Groen et al., 2022; Kay et al., 2015)***. Critically, the slope of the line relating pRF eccentricity to pRF size increases as one ascends through the visual hierarchy. The previous results – highly similar mean pRF size between DPs and controls – could therefore have hidden atypical scaling of pRF size and eccentricity. Therefore, to compare how pRF sizes scale with eccentricity within DPs and controls, ROI-specific LMMs were created that modeled the linear relationship between pRF size and pRF eccentricity with subject-specific intercepts and slopes as random effects. Then, the effects of group, hemisphere, and their interaction were introduced as fixed effects. Parameter estimates from each ROI’s model are displayed in Table 1 and subject-specific slopes by group and ROI are displayed graphically in Figure 3. Since multiple models were constructed, a conservative alpha, *α* < 0.01, was adopted to guide statistical interpretation. In none of the ROIs did group membership, hemisphere, or the interaction between the two significantly explain additional variance in relating pRF eccentricity to pRF size (all *ps* > 0.01). Hence, DPs and controls show highly similar relationships between pRF size and pRF eccentricity.

**Figure 3.**
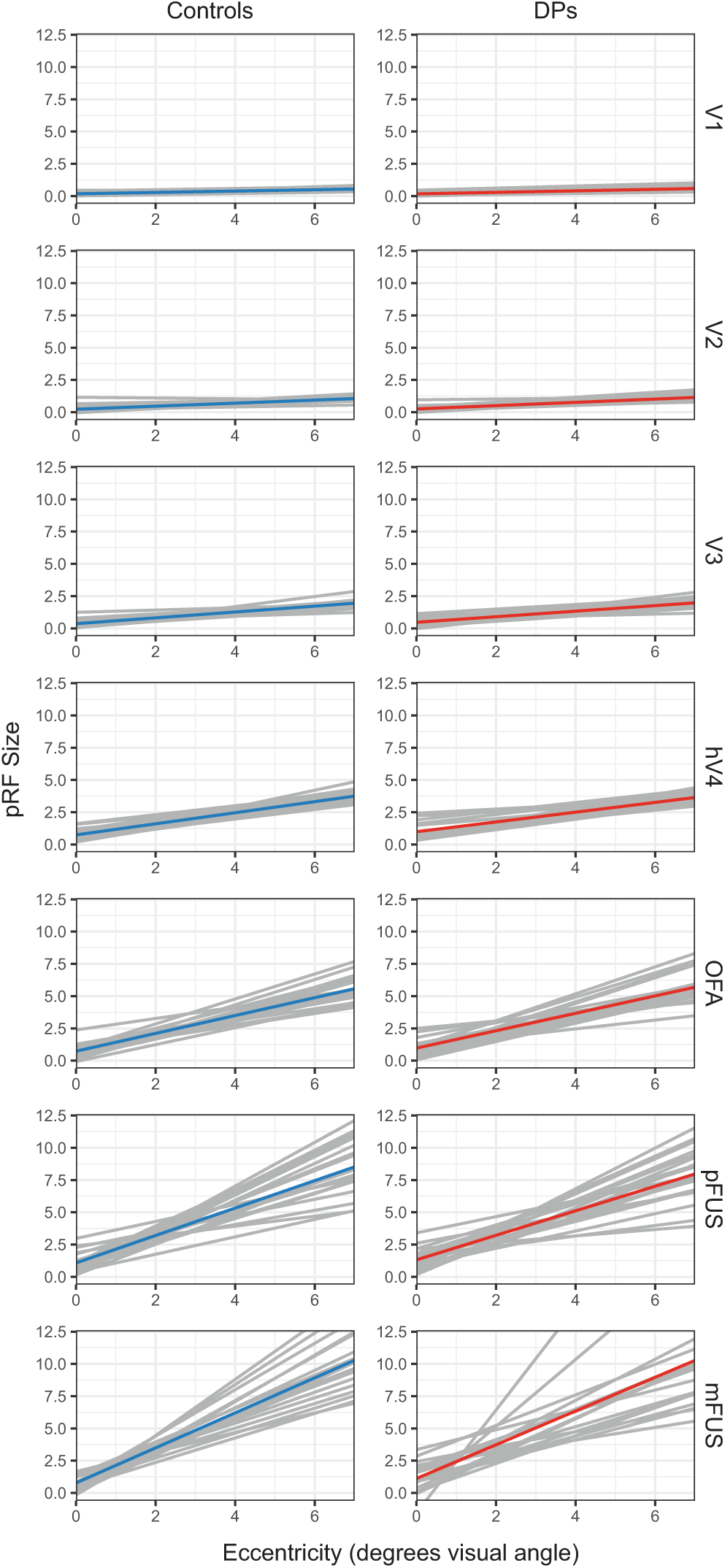
Lines of best fit relating pRF eccentricities (in degrees of visual angle) to pRF sizes by group and ROI. Gray lines show lines of best fit for individual subjects and colored lines show the group average. Lines were constrained to the central 6 degrees of visual angle because data in far eccentricities were often sparse, especially in more anterior face-selective regions.

**Table 1.**
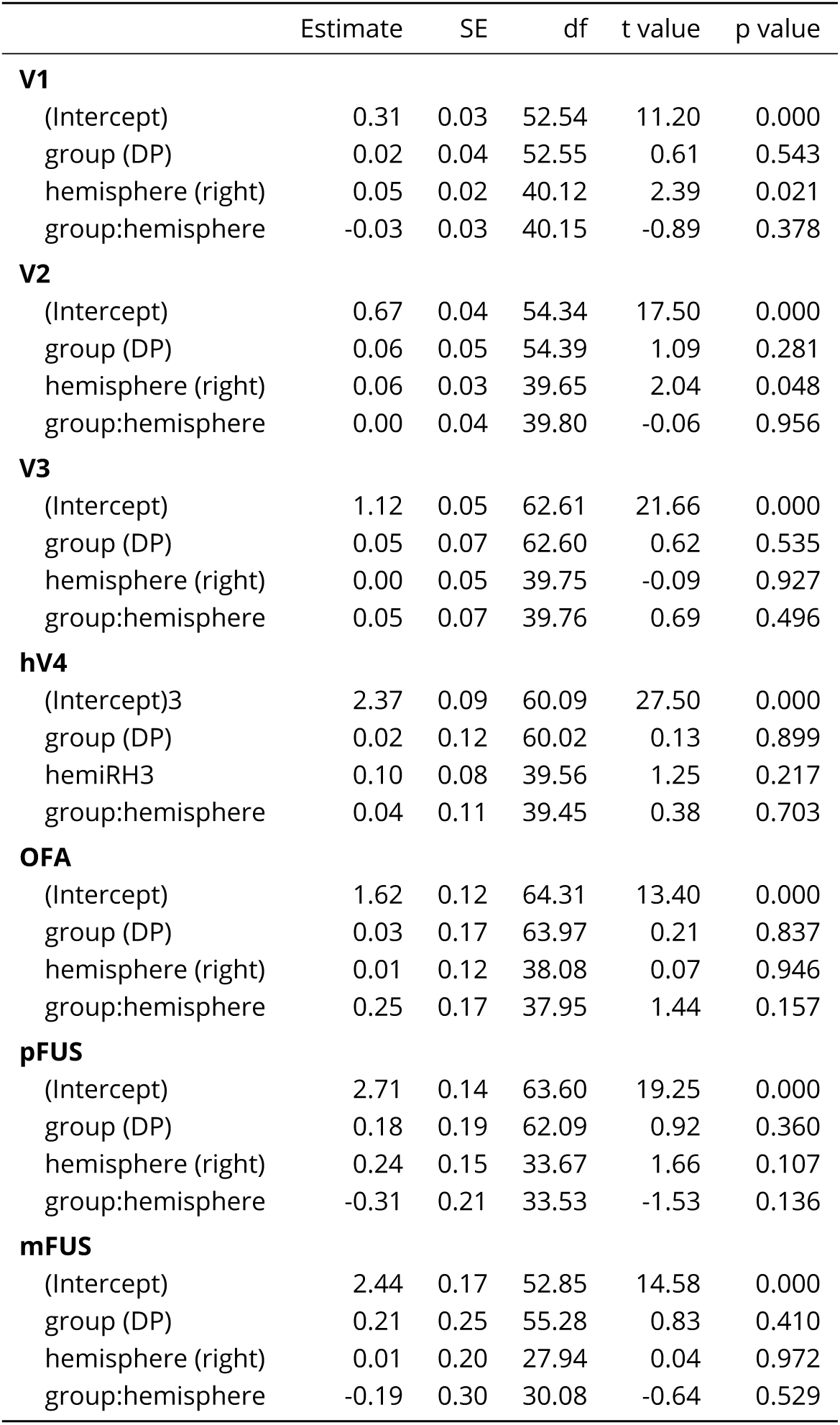
Fixed effect parameter estimates of pRF size by hemisphere (right, left) for the control and developmental prosopagnosic (DP) groups. Separate models were created for each region of interest. Note that in each one the intercept has been mapped to the control group in the left hemisphere. Formula (R, lme4 package): size ~ group + hemisphere + group:hemisphere + (1+eccentricity|subjectID) ∗ *p* < 0.01; ∗∗ *p* < 0.001

|  | Estimate | SE | df | t value | p value |
| --- | --- | --- | --- | --- | --- |
| **V1** |  |  |  |  |  |
| (Intercept) | 0.31 | 0.03 | 52.54 | 11.20 | 0.000 |
| group (DP) | 0.02 | 0.04 | 52.55 | 0.61 | 0.543 |
| hemisphere (right) | 0.05 | 0.02 | 40.12 | 2.39 | 0.021 |
| group:hemisphere | -0.03 | 0.03 | 40.15 | -0.89 | 0.378 |
| **V2** |  |  |  |  |  |
| (Intercept) | 0.67 | 0.04 | 54.34 | 17.50 | 0.000 |
| group (DP) | 0.06 | 0.05 | 54.39 | 1.09 | 0.281 |
| hemisphere (right) | 0.06 | 0.03 | 39.65 | 2.04 | 0.048 |
| group:hemisphere | 0.00 | 0.04 | 39.80 | -0.06 | 0.956 |
| **V3** |  |  |  |  |  |
| (Intercept) | 1.12 | 0.05 | 62.61 | 21.66 | 0.000 |
| group (DP) | 0.05 | 0.07 | 62.60 | 0.62 | 0.535 |
| hemisphere (right) | 0.00 | 0.05 | 39.75 | -0.09 | 0.927 |
| group:hemisphere | 0.05 | 0.07 | 39.76 | 0.69 | 0.496 |
| **hV4** |  |  |  |  |  |
| (Intercept)3 | 2.37 | 0.09 | 60.09 | 27.50 | 0.000 |
| group (DP) | 0.02 | 0.12 | 60.02 | 0.13 | 0.899 |
| hemiRH3 | 0.10 | 0.08 | 39.56 | 1.25 | 0.217 |
| group:hemisphere | 0.04 | 0.11 | 39.45 | 0.38 | 0.703 |
| **OFA** |  |  |  |  |  |
| (Intercept) | 1.62 | 0.12 | 64.31 | 13.40 | 0.000 |
| group (DP) | 0.03 | 0.17 | 63.97 | 0.21 | 0.837 |
| hemisphere (right) | 0.01 | 0.12 | 38.08 | 0.07 | 0.946 |
| group:hemisphere | 0.25 | 0.17 | 37.95 | 1.44 | 0.157 |
| **pFUS** |  |  |  |  |  |
| (Intercept) | 2.71 | 0.14 | 63.60 | 19.25 | 0.000 |
| group (DP) | 0.18 | 0.19 | 62.09 | 0.92 | 0.360 |
| hemisphere (right) | 0.24 | 0.15 | 33.67 | 1.66 | 0.107 |
| group:hemisphere | -0.31 | 0.21 | 33.53 | -1.53 | 0.136 |
| **mFUS** |  |  |  |  |  |
| (Intercept) | 2.44 | 0.17 | 52.85 | 14.58 | 0.000 |
| group (DP) | 0.21 | 0.25 | 55.28 | 0.83 | 0.410 |
| hemisphere (right) | 0.01 | 0.20 | 27.94 | 0.04 | 0.972 |
| group:hemisphere | -0.19 | 0.30 | 30.08 | -0.64 | 0.529 |

### DPs have normal distributions of pRF centers

Our results show that DPs and controls have very similar VFC. However, since VFC depends on both pRF sizes and pRF center locations – with larger pRFs covering more visual space and pRFs typically increasing in size with eccentricity – similar VFC patterns can arise even if the groups differ in the underlying spatial distribution of pRF centers. In other words, if one group had pRFs located more peripherally, the accompanying increase in pRF size could lead to comparable VFC between the groups despite underlying differences in where the pRFs are located. Therefore, in the next analysis, we directly compare groups on the distributions of pRF center locations.

To this end, we computed the area of the 95% probability contour encircling the bivariate distributions of *x* and *y* coordinates of pRF centers. Larger values indicate greater dispersion of pRF centers. In none of the ROIs (V1-hV4, OFA, pFUS, or mFUS) did the pRF dispersion differ significantly between DPs and controls (two sample *t*-tests, uncorrected, all *ps* > 0.05). The group-wise density of pRF center locations across all ROIs is represented in Figure 4A, and they show similar patterns between groups. Both groups also showed similar proportions of total pRFs occupying distinct eccentricity bins (Figure 4 B).

**Figure 4.**
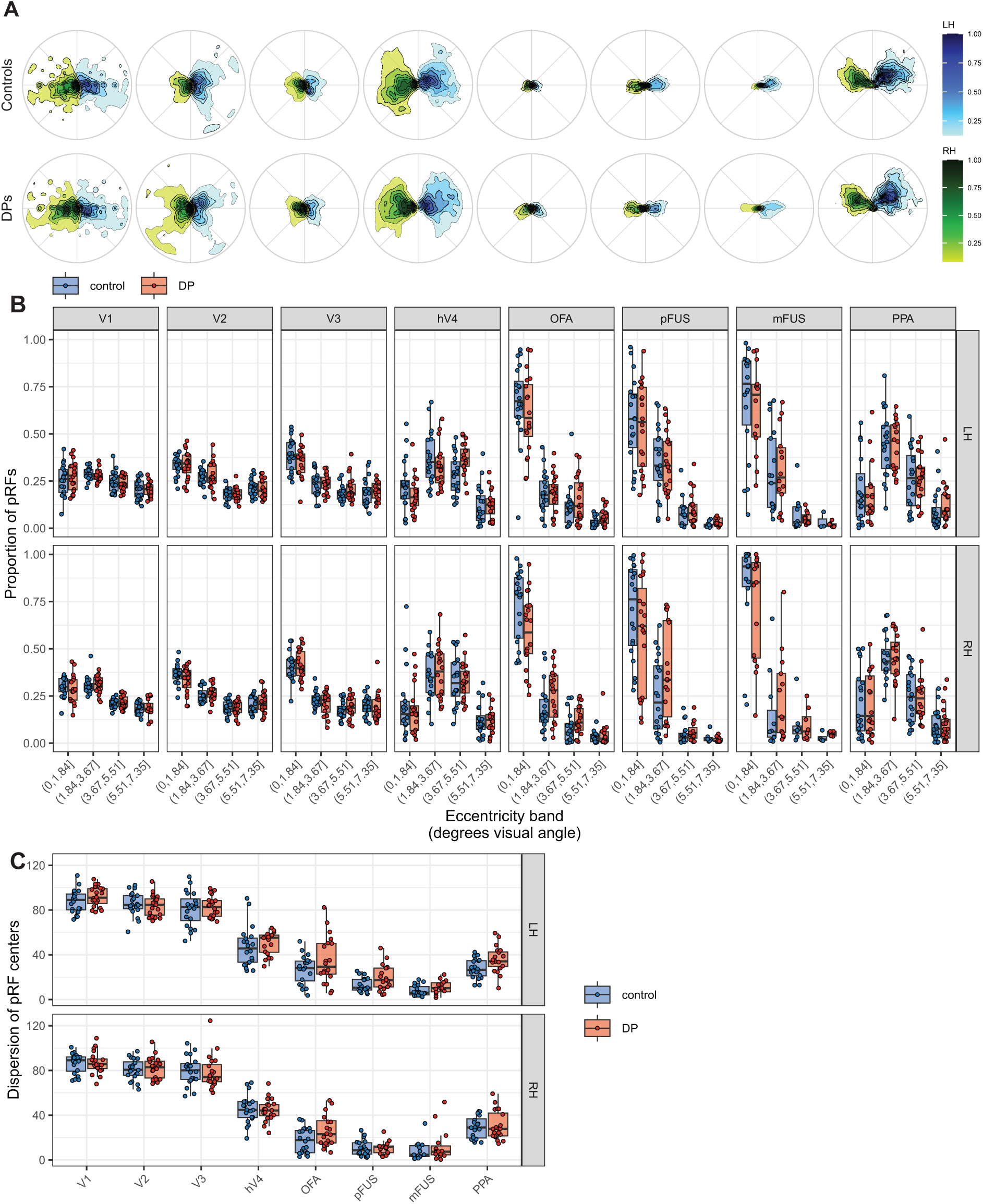
Spatial distributions of pRF center coordinates across ROIs and hemispheres. A) Contours representing the group density distributions of pRF center coordinates by group and hemisphere. Green colors represent pRFs in the right hemisphere and blue colors represent pRFs in the left hemisphere. B) The proportion of pRFs falling into four distinct eccentricity bins. Datapoints represent individual participants. C) Measures of bivariate dispersion of pRF centers by group and ROI. Datapoints represent individual participants. Bivariate spread was measured by computing the area of the 95% probability contour encircling the distribution of pRF centers.

Finally, to rigorously evaluate our claim that spatial integration is ‘normal’ in DP, we conducted Bayesian *t*-tests comparing mean pRF size between groups (Supplementary Table 6). Using a default Cauchy prior on effect size (rscale = medium), evidence consistently favored the null hypothesis in 15 out of 16 regions (*BF*_01_ range 1.25 to 3.16). The right OFA was the only region showing moderate evidence for a group difference (*BF*_10_ = 3.85). Crucially, however, pRFs in this region were actually larger in DPs than in controls (DPs: mean pRF size ± SD = 2.42±0.82; Controls: 1.78±0.75). Because the prevailing theory explicitly posits that *abnormally small* pRFs underlie DP ***(Avidan and Behrmann, 2021; Witthoft et al., 2016)***, this deviation provides no support for the restricted spatial integration hypothesis. Furthermore, when we adjusted the prior to an ‘ultrawide’ scale to reflect the massive effect size reported by ***Witthoft et al. (2016)***, evidence heavily favored the null hypothesis across the visual hierarchy (i.e. *BF*_01_ range 2.46 to 5.00 in all face-selective regions except right OFA). Therefore, our data confidently reject the presence of the severe pRF size reductions previously thought to underlie DP.

In summary, across multiple measures of retinotopic organization, DPs show no deviation from control participants. DPs and controls have highly similar coverage of the visual field, scaling between pRF eccentricity and size, and distribution of pRF centers.

## Discussion

Population receptive field (pRF) mapping is a valuable technique in visual neuroscience that has provided an understanding of the location, size, and shape of aggregate receptive fields across the cortex ***(Groen et al., 2022; Wandell and Winawer, 2015)***, however few results have demonstrated strong relationships between individual variation in pRF properties and behavioral measures of perception. One of the most striking studies showing a link between pRF properties and behavior reported that a group of individuals with developmental prosopagnosia (DP) had exceptionally small and foveally-concentrated pRFs in intermediate visual areas and face-selective regions ***(Witthoft et al., 2016)***. Moreover, pRF size correlated with their face recognition performance. These findings presented an appealing neural explanation for the diminished holistic face processing sometimes observed in DPs ***(Avidan and Behrmann, 2021; Avidan et al., 2011; Degutis, 2012; Palermo et al., 2011)***.

To rigorously investigate spatial integration in DPs, we used fMRI to measure pRF properties in DPs and controls (*N* = 20 per group) using high-level mapping stimuli capable of robustly stimulating both early and category-selective visual regions. In contrast to ***Witthoft et al. (2016)***, our results provide strong empirical evidence that pRF properties are *highly similar* between DPs and controls. Both groups showed comparable: 1) visual field coverage (VFC) maps, providing a measure of how pRFs within an ROI collectively tile the visual field; 2) scaling between pRF eccentricity and pRF size, both within regions of interest (ROIs) and across the visual hierarchy; and 3) dispersion of pRF centers around the visual field. These three patterns of results held for early, intermediate, and ventral face-selective areas, as well as a category-selective control ROI not involved in face processing (the scene-selective parahippocampal place area or PPA; ***Epstein and Kanwisher (1998)***). Similarly, we found no significant correlations between behavioral face recognition ability (assessed by either the CFMT or Famous Faces Test) and any of the main pRF measures surveyed here, including pRF size, eccentricity, VFC area, and dispersion of centers (for full statistical reporting, see Supplementary Table 5).

### Methodological strengths of the current study

Our methods incorporated several features that ensure the validity and robustness of the findings. First, we employed a carefully thought-out, ecologically valid stimulus capable of significantly stimulating both early visual and category-selective regions. The stimulus featured vivid, high-contrast content including images of faces, objects, bodies, and places randomly placed on a background of pink noise. Although high-level regions have been mapped with low-level stimuli before (e.g., black– and-white checkerboard patterns), these types of stimuli do not drive category-selective regions as strongly, producing poorer goodness-of-fit ***(Le et al., 2017; Silson et al., 2015)*** and unstable pRF estimates.

Second, we acquired in-scanner eye-tracking data during the retinotopic mapping sessions whenever it was feasible. Eye movements can severely distort pRF measurements, usually by increasing estimated pRF size, so it was critical to compare fixation behavior in the DPs and controls. Participants with usable eye-tracking data demonstrated steady fixation, with no significant differences in gaze stability between DPs and controls. For participants without usable eye tracking data – due to either interference from thick prescription lenses or incompatibility of head anatomy and hardware setup – consistent fixation was confirmed by checking for clearly delineated polar angle and eccentricity maps in early visual cortex (Supplemental Figures 6 and 7).

One explanation for the comparable results for the DPs and controls is that the putative DPs may not have possessed a true face recognition deficit. However, we are confident that our selection process effectively categorized DPs for the following reasons. First, all DPs self-reported substantial face recognition difficulties affecting their daily lives, quantified by scores on the PI-20 questionnaire ***(Shah et al., 2015)***. Furthermore, objective inclusion criteria required DPs to score more than 1.8 standard deviations below the mean of a control sample on two independent tests of face recognition – the CFMT and a famous faces test – a stringent threshold consistent with or exceeding those used in numerous studies of DP ***(DeGutis et al., 2023)***. Furthermore, using not one but two tests for classification substantially minimizes the risk of false identification. Additional testing with the DPs was conducted to rule out low-level visual deficits or autistic-like traits as alternative causes for poor face performance. Univariate measures of face-selectivity provided another opportunity to confirm that our DP participants showed the neural abnormalities that have been found in previous fMRI studies of DP ***(Furl et al., 2011; Jiahui et al., 2018)***. Univariate functional localizer analyses from our pool of participants revealed that, at the group level, DPs compared to controls exhibited significantly lower face-selective activation in many core ventral face-selective regions (See Supplemental Table 3), replicating previous findings ***(Dinkelacker et al., 2011; Furl et al., 2011; Gerlach et al., 2019; Jiahui et al., 2018)***. Notably, limiting voxels to only those that showed strong retinotopic modulation, still revealed significant reductions in face-selectivity in DPs (Figure 1 B) a novel finding that provides additional evidence that DPs were classified correctly.

Confidence in our pRF mapping methods was further supported by the fact that our results replicated many key pRF phenomenon from prior studies. Both groups showed significant increases in pRF size across the visual processing hierarchy ***(Finzi et al., 2021; Kay et al., 2015; Silson et al., 2015)*** and, within ROIs, there was a consistent increase in pRF size with pRF eccentricity ***(Benson et al., 2018; Kay et al., 2015)***. Regardless of group, the three ventral face-selective areas exhibited a higher proportion of foveally-positioned pRFs than early visual areas ***(Grill-Spector et al., 2018; Kay et al., 2015; Silson et al., 2015)***. Furthermore, these face-selective areas showed a relatively higher degree of subadditive spatial summation (larger static nonlinearity parameter in the CSS model; ***Kay et al., 2015***, ***2013a***,b). Also, we replicated previously reported visual field biases – namely, a lower visual field bias in OFA ***(Gomez et al., 2018; Silson et al., 2015)*** and an upper visual field bias in PPA ***(Silson et al., 2015)***. Across both groups, qualitative inspection of polar angle maps near the calcarine sulcus in each hemisphere revealed the canonical retinotopic organization of smooth sweeps from the upper vertical meridian to the lower vertical meridian through the contralateral side of the visual field. Eccentricity maps likewise showed the expected progression from foveal pRFs near the occipital pole to increasingly more peripheral pRFs anteriorly and medially ***(Benson et al., 2018)***. Together, these multiple converging lines of evidence confirm that data quality was high and closely matched between groups.

### How do our pRF results fit in with other behavioral data from DPs?

Our results directly challenge the view that DP stems from impaired spatial integration caused by abnormally small pRFs. While this view has garnered substantial interest ***(Avidan and Behrmann, 2021; Grill-Spector et al., 2018; Witthoft et al., 2016)***, some behavioral evidence is inconsistent with it. Behaviorally, the most common tasks used to evaluate holistic face processing are the composite ***(Young et al., 1987)*** and part-whole tasks ***(Tanaka and Farah, 1993)***. Comparisons between DPs and controls on these measures have yielded some mixed results: some report reduced holistic processing ***(Avidan et al., 2011; Degutis, 2012; Palermo et al., 2011)***, while others find it intact ***(Biotti et al., 2017; Le Grand et al., 2006; Susilo et al., 2010)***. This inconsistency highlights the heterogeneous nature of DP and complicates linking DP to a single underlying cause such as poor spatial integration.

Behavioral face studies that experimentally restrict spatial integration further challenge the notion that spatial integration deficits underlie DP. For instance, ***Tsantani et al. (2020)*** compared the ability of DPs and controls to identify famous faces under two viewing conditions: when faces were presented briefly in their entirety versus progressively revealed by a dynamic, narrow aperture. The dynamic aperture viewing condition allowed participants to inspect local features of the face but prevented distant regions from being processed simultaneously, thus simulating the effect of having a small window of integration. In this condition, control participants exhibited substantial decrements in performance likely because it hindered holistic face processing mechanisms. If the DPs’ deficits at face recognition stemmed from deficient sampling of the visual field by pRFs, one would expect the dynamic aperture viewing condition to elicit a smaller decrement in performance. However, contrary to this prediction, both groups showed comparable performance decrements. Similar results were found when DPs were tested with gaze-contingent displays restricting information to a small region around the fovea ***(Verfaillie et al., 2014)***, reinforcing the conclusion that spatial integration deficits alone do not underlie DP.

Lastly, DP is defined by deficits with facial identity processing. However, if DP is a deficit in spatial integration, individuals with DP should have difficulties with *any* aspect of face processing that involves holistic perception. Facial sex judgments ***(Baudouin and Humphreys, 2006)*** and facial expression recognition ***(Calder et al., 2000)*** both elicit composite effects that are similar in size to composite tasks involving facial identity recognition. Yet, most DPs perform as well as controls on non-identity face perception tasks, such as sex classification ***(Chatterjee and Nakayama, 2012; Degutis, 2012; Marsh et al., 2019)*** and facial expression recognition ***(Bell et al., 2023; Palermo et al., 2011)***. So, DPs do not appear to have broad deficits with spatial integration of facial information ***(Baudouin and Humphreys, 2006; Calder et al., 2000; Durand et al., 2007; Yokoyama et al., 2014)***.

### What factors may have contributed to the different results for the present study and Witthoft et al. (2016)?

In light of the difference in findings between our study and that of ***Witthoft et al. (2016)***, several methodological differences between the two studies are worth considering. First, while ***Witthoft et al. (2016)*** mapped pRFs using black-and-white checkerboard patterns as the carrier image, we employed a more ecologically valid stimulus featuring colorful high-level images including faces, scenes, body parts, and objects. Although both types of stimuli would be expected to robustly stimulate early visual cortex, our stimulus better targets higher-level regions as reflected in noticeably higher goodness of fit, *R*^2^, in our data compared to that of ***Witthoft et al. (2016)*** (current study mean *R*^2^: OFA=.45, pFUS=.42, mFUS=.33; ***Witthoft et al. (2016)*** approximate mean *R*^2^: OFA=.21, pFUS=.15, mFUS=.07;). This boost agrees with prior within-subject studies demonstrating that more naturalistic and region-appropriate mapping stimuli outperform black-and-white checkerboard patterns in category-selective areas ***(Le et al., 2017; Silson et al., 2015)***.

More critically, the choice of stimulus can systematically alter pRF parameters themselves. For instance, ***Le et al. (2017)*** showed that visual field coverage (VFC) in ventral occipito-temporal cortex, a word form-selective region, shrinks toward the central visual field when mapped with word stimuli compared to checkerboard patterns. Thus, the results with DPs reduced VFC with checker-boards but normal VFC with more naturalistic mapping stimuli – may reflect a stimulus-dependent effect that is elicited solely by low-level, non-preferred stimuli in face-selective regions. Nevertheless, our findings, demonstrating normal spatial integration in face-selective regions when using more appropriate, naturalistic stimuli offer a more definite and convincing proof of neural tuning in these higher-level regions.

To model pRFs, we used the compressive spatial summation (CSS; ***Kay et al. (2013a***,b)) model as opposed to a linear pRF model as used in ***Witthoft et al. (2016)***. The main difference between these two models is that the CSS model applies a compressive static nonlinearity to account for subadditive spatial summation (situations where the sum of the individual responses to apertures forming complementary pairs is < 1). Subadditive spatial summation is present in EVC and grows more pronounced in higher-order regions, making this model more appropriate for studying spatial integration in face-selective regions. Both groups, however, evinced highly similar values for the compressive static nonlinearity parameter across the ROIs studied (Figure 1G). Because this nonlinear exponent parameter did not considerably differ between DPs and controls, the use of a linear model in prior work ***(Witthoft et al., 2016)*** is unlikely to explain the massive group differences they reported, as it would be expected to scale pRF estimates similarly for both groups.

It is also worth noting that ***Witthoft et al. (2016)*** mapped pRFs with stimuli spanning approximately 30° in diameter, compared to 14.7° in the present study, a substantial difference in field of view. For us, it was necessary to limit the field of view presented inside the scanner in order to collect high-quality eye-tracking data for verification of stable fixation, which we consider a critical step. Nevertheless, the field of view we used still greatly exceeds the typical visual angle subtended by a face at normal conversational distance (~8°, ***McKone (2009)***), so this design appears to provide a sufficiently wide enough visual field to detect any spatial integration deficits in DPs, if present.

Finally, there were differences in the aperture design used to map pRFs. ***Witthoft et al. (2016)*** employed blocked runs of rotating wedges and expanding rings (2 runs each, 4 runs total), whereas we used a sequence of apertures including bars (sweeping in 8 different directions), one rotating wedge, and a brief full-field stimulation. While the precise impact of aperture design on pRF estimates is not well understood (for one investigation see ***Alvarez et al. (2015)***), our aperture design successfully replicated many established pRF properties and it remains unclear why this difference between the designs would have differentially affected DPs and controls. Moreover, our study collected more than twice as much data per participant (5 runs of 6.8 mins each versus 4 runs of 3.7 mins each), enhancing the precision and reliability of pRF parameter estimates, especially when combined with a more effective mapping stimulus.

## Conclusion

Although we agree that abnormally small pRFs and restricted VFC in face-selective regions could cause deficits to mechanisms supporting face recognition, our results – using rigorous pRF mapping methods, a large sample, and region-appropriate mapping stimuli – indicate that spatial integration, as measured by pRFs, is intact in DPs. Therefore, the pRF processing framework, while useful in conceptualizing many aspects of visual function, does not appear to provide insight into the cognitive and neural deficits underlying DP. This underscores the need for a more nuanced and multi-faceted understanding of the computations that underlie face processing. Many different factors, besides visual spatial processing by pRFs, are likely to contribute to effective face representations ***(Kay and Yeatman, 2017)***, and the various forms of DP may arise from abnormalities in one or a combination of these. For instance, models may need to incorporate specific aspects of neural tuning to facial information, such as sensitivity to face parts and their relational configurations ***(Arcaro et al., 2020; Poltoratski et al., 2021)***. This would mark a shift towards a more general characterization involving representational content rather than receptive field positioning. In other cases, the disorder may be explainable through abnormalities in functional connectivity ***(Avidan et al., 2014; Lohse et al., 2016; Rosenthal et al., 2017)*** or structural connectivity ***(Gomez et al., 2015; Song et al., 2015b)***. One observation, though, that appears robust at the group level is diminished face selectivity in DP, which may serve as an anchor for future investigations. Given the heterogeneity likely present in DP, identification of the distinct mechanisms that contribute to face recognition impairments may therefore require behavioral and neuroimaging methods capable of characterizing deficits at the level of individual participants.

## Materials and Methods

### Participants

Twenty individuals with DP (15 females, *M_age_* = 38.18, *SD_age_* = 11.13) and 20 typical controls (14 females, *M_age_* = 33.86, *SD_age_* = 10.92) participated in the study. The mean age of the groups did not differ significantly (*t*(40.97) = 1.286, *p* = .206, *d* = 0.398, *CI*_95%_ = –0.216, 1.006). Written informed consent was obtained from all participants in accordance with the Declaration of Helsinki and a protocol approved by the Dartmouth College Committee for the Protection of Human Subjects (#23282).

### Diagnostic testing and participant selection

DP participants were recruited from our prosopagnosia database (www.faceblind.org), and interested participants first completed the Twenty-Item Prosopagnosia Index (PI20; ***Shah et al. (2015)***). The PI20 responses from every DP participant enrolled in the study indicated substantial face recognition difficulties that affected their daily lives (M = 81.90/100, SD = 8.86).

Face recognition ability was assessed with two online tests: The Cambridge Face Memory Test (CFMT; ***Duchaine and Nakayama (2006)***) and a Famous Faces Test (FFT; ***Kieseler and Duchaine (2023)***). Individual scores and control summary statistics are presented in Table 2. The famous faces test was newly created and required participants to identify (via free response) the faces of 40 celebrities familiar to North Americans. Immediately after completing the test, participants were asked about their familiarity with each celebrity. Human raters judged if each response was sufficiently correct, and each participant’s performance was quantified as the number of correct responses out of the total number of celebrities they reported being familiar with. Potential DPs’ face recognition was deemed sufficiently impaired if they scored more than 1.8 standard deviations below the control sample means on both the CFMT and FFT. Scores on the CFMT were compared against data from 50 typical observers reported by ***(Duchaine and Nakayama, 2006)***, and scores on the FFT were compared against data from N typical observers collected online through www.testable.org.

**Table 2.** Scores from each member of the prosopagnosic sample on diagnostic tests consisting of: The Hanover Early Visual Assessment (HEVA), The 20-Item Prosopagnosia Index (PI20), The Cambridge Face Memory Test (CFMT), The Cambridge Face Memory Test with Australian Faces (CFMT Aus), famous faces test (FFT), old/new faces test, and The Subthreshold Autism Trait Questionairre (SATQ). Asterisks (*) indicates a given score meets the inclusion criterion set in advance of the study.

| identifier | Sex | Age | HEVA<br>max: 120 | PI20<br>max: 100 | Face processing diagnostic tests |  |  |  | SATQ<br>max: 57 |
| --- | --- | --- | --- | --- | --- | --- | --- | --- | --- |
|  |  |  |  |  | CFMT<br>max: 72 | CFMT Aus<br>max: 72 | FFT<br>correct/familiar | Old/New Faces<br>(A Prime) |  |
| sub-2432 | female | 28 | 101 | 67 | 36* | - | 16/31=0.52* | - | 19* |
| sub-2455 | female | 48 | 84 | 82 | 42* | - | 25/38=0.66* | - | 14* |
| sub-2507 | female | 30 | 98 | 86 | 39* | - | 15/29=0.52* | - | 7* |
| sub-2538 | male | 33 | 104 | 86 | 39* | - | 16/36=0.44* | - | 2* |
| sub-2550 | female | 55 | 107 | 84 | 40* | - | 14/28=0.5* | - | 11* |
| sub-2582 | male | 31 | 88 | 94 | 36* | - | 8/28=0.29* | - | 3* |
| sub-2645 | female | 26 | 102 | 79 | 37* | - | 10/21=0.48* | - | 16* |
| sub-2671 | female | 44 | 100 | 90 | 43* | - | 23/38=0.61* | - | 16* |
| sub-2687 | female | 52 | 100 | 62 | 43* | - | 18/34=0.53* | - | 16* |
| sub-2699 | male | 24 | 76 | 80 | 39* | - | 12/27=0.44* | - | 12* |
| sub-2709 | female | 41 | 97 | 81 | 44 | 40* | 11/40=0.28* | - | 8* |
| sub-2733 | female | 53 | 88 | 79 | 25* | - | 29/40=0.72 | 0.4* | 22* |
| sub-2739 | female | 23 | 101 | 94 | 39* | - | 4/27=0.15* | - | 16* |
| sub-2750 | male | 47 | 102 | 74 | 31* | - | 4/27=0.15* | - | 24* |
| sub-2779 | female | 53 | 102 | 73 | 41* | - | 3/25=0.12* | - | 13* |
| sub-2804 | female | 28 | 96 | 94 | 37* | - | 9/36=0.25* | - | 14* |
| sub-2820 | male | 29 | 90 | 93 | 40* | - | 17/39=0.44* | - | 26* |
| sub-2828 | male | 40 | 95 | 79 | 42* | - | 19/39=0.49* | - | 12* |
| sub-2859 | female | 24 | 89 | 82 | 34* | - | 22/34=0.65 | 0.67* | 10* |
| sub-2935 | female | 54 | 98 | 81 | 37* | - | 21/39=0.54* | - | 22* |
| DP Mean: |  | 38.15 | (93.4)<sup>1</sup>(97.9)<sup>2</sup> | 82.00 | 38.20 | - | (0.42)<sup>3</sup>(0.46)<sup>4</sup> | - | 14.15 |
| DP SD: |  | 11.70 | (8.52)<sup>1</sup>(6.80)<sup>2</sup> | 8.75 | 4.49 | - | (0.15)<sup>3</sup>(0.21)<sup>4</sup> | - | 6.47 |
| Control Mean: |  | 33.86 | (93.6)<sup>1</sup>(86.4)<sup>2</sup> | - | 58 | 57.8 | (0.86)<sup>3</sup>(0.91)<sup>4</sup> | 0.96 | 15.87 |
| Control SD: |  | 10.92 | (11.0)<sup>1</sup>(15.5)<sup>2</sup> | - | 7.9 | 7.82 | (0.17)<sup>3</sup>(0.11)<sup>4</sup> | 0.02 | 6.89 |
| Inclusion criterion: |  | - | Z < -1.8 | - | Z < -1.8 | Z < -1.8 | Z < -1.8 | Z < -2 | Z < 2 |

In cases where potential DP participants met the inclusion criteria for only one of the two face recognition tests, participants were given a third test. If a participant’s CFMT score was in the normal range, they were administered the CFMT-Australian ***(McKone et al., 2011)***; if their Famous Face Test score was in the normal range, they were tested with an old/new test involving female faces ***(Duchaine and Nakayama, 2005)***. The CFMT-Australian follows the same format and scoring as the original CFMT. The old/new test required memorization of ten target faces, followed by 50 test trials (20 targets plus 30 distractors). Performance on the old/new test was evaluated by calculating A prime scores. Participants were included in the study if they scored below the threshold (CFMT-Australian: *Z* < –1.8; Old/new faces: *Z* < –2) on either of the follow-up tests.

To ensure that the DPs’ impairments in face recognition reflected a high-level deficit rather than a deficit to early visual processing, each DP recruit was assessed with the Hanover Early Visual Assessment ***(Kieseler et al., 2022)***. Any potential recruit with a score below 1.8 SDs below the mean of a control sample (*N* = 117) was not given further consideration for the study. This left a final sample of DPs that scored very similarly to controls (Controls: mean±SE = 90.0±13.3; DPs: mean±SE = 95.6±7.8).

Because rates of face recognition impairments are elevated in Autism Spectrum condition ***(Griffin et al., 2021)***, participants were also screened for subthreshold autistic traits using the self-report Subthreshold Autism Trait Questionnaire (SATQ; ***Kanne et al. (2012)***). In our analysis of the SATQ responses, we omitted 5 of the 24 questions that measured behavioral tendencies likely to be influenced by poor face recognition (“*I enjoy social situations where I can meet new people and chat (i.e. parties, dances, sports, and games)*”; “*I seek out and approach others for social interactions*”; “*Others think I am strange or bizarre*”; “*I have some behaviors that others consider strange or odd*”; “*I make eye contact when talking with others*”). All DPs were less than 2 SDs above the mean of a control sample (*N* = 133; ***Bell et al. (2023)***).

### Experiment design

***Session 1: Functional localizer.*** To localize face-selective regions of interest (ROIs), a category localizer was run that presented six classes of stimuli: faces, bodies, objects, phase-scrambled objects, natural scenes, and words. Stimuli consisted of dynamic video clips featuring various visual elements in motion, all filmed against a uniform black backdrop to maximize contrast and visibility. Videos of the natural scenes filled the entire frame and displayed a slow forward camera movement equivalent to a normal walking pace from a first-person perspective. Video clips subtended ~26° *x* 15° of visual angle in width and height. Participants viewed four runs, each of which lasted 8 minutes and 22 seconds. Within each run, stimuli were grouped into 14-second blocks containing five videos each. Blocks of each category were displayed four times in each run in a quasi-random order, twice in color and twice in grayscale. The trial order was the same for all participants. Throughout the experiment, participants were allowed to freely move their eyes and instructed to identify back-to-back repeats with a button press (1-back task). Each of the four runs presented unique stimuli.

Face-selective ROIs were created by analyzing voxels within masks centered on associated anatomical landmarks and selecting the 20% of the voxels within the mask with the highest *t*-values for the contrast comparing faces versus objects. This method was chosen for its ability to generate individually tailored functional ROIs, while at the same time ensuring equal voxel counts across participants as well as retention of data from participants whose activations might otherwise fall below fixed thresholds ***(Jiahui et al., 2018; Norman-Haignere et al., 2013)***. Additionally, an ROI on the parahippocampal gyrus selective more for scenes than objects (parahippocampal place area or PPA ***Epstein and Kanwisher, 1998)*** was included to serve as a category-selective control region not involved in face perception.

***Session 2: pRF mapping.*** Each participant completed five runs of retinotopy to identify the portion of the visual field capable of eliciting a response from the population of neurons within a voxel (i.e., each voxel’s pRF). In each run, participants fixated a central dot (0.2° x 0.2°) while apertures of various shapes and sizes gradually swept across the visual field revealing colorful high-level images flickering at a rate of 5 Hz.

To elicit a strong BOLD response across a variety of category-selective areas, we used stimuli introduced by ***Benson et al. (2018)***. These stimuli were created by taking cutout images from ***Kriegeskorte et al. (2008)*** (containing faces, body parts, foods, objects, street signs, animals, and architectural elements, etc.) and placing them at random positions and scales on achromatic pinknoise backgrounds (1/*f* amplitude spectrum). As such, we refer to them as *“image mashup”*. Our early pilot studies demonstrated the image mashup stimuli were effective at driving responses in face-selective regions but not scene-selective regions, which were of tangential interest to our study of DP (see ***Jiahui et al. (2018)***). Therefore, we interleaved the image mashup stimuli with photographs of natural landscapes, and pilot data showed this combined approach generated a strong response in both face-selective *and* scene-selective regions. The image mashup and natural landscape stimuli alternated at a rate of 5 Hz, so each 2 s aperture presentation included an equal number of presentations of each stimulus type. Apertures and images were prepared at a resolution of 1060 x 1060 pixels and were constrained to a central circular region of diameter 14.7°. All stimuli were presented against a medium gray background with a faint semi-transparent polar grid pattern superimposed to facilitate fixation.

Three different types of apertures were presented in every run – a full field disk, a sequence of bars, and a rotating wedge (a schematic diagram is included in Supplemental Figure 5). Apertures progressed the same way in every run: After 8 s of initial blank fixation, a disk-shaped aperture appeared that revealed stimuli within the entire circular extent of the visual field, lasting for a duration of 4 s. Following 6 s of rest, bar-shaped apertures traversed the visual field in sweeps of 18 discrete, evenly spaced steps, each lasting 2 s. Each bar presentation overlapped 50% of the area of the previous position. Eight sweeps occurred, each interleaved with 6 s of rest, in the following order for all runs: Top–Bottom, Bottom–Top, Right–Left, Left–Right, Upper Right–Bottom Left, Bottom Left–Upper Right, Upper Left–Bottom Right, Bottom Right–Upper Left. Following another 6 s of rest, a 90° wedge appeared in the upper left quadrant and made 16 discrete clockwise steps, 2 s each, sweeping a full 360° around the central circular visual field (22.5° turn per step). Each run ended with 17 s of final fixation.

Throughout all retinotopy scans, participants performed a color detection task at fixation in which they reported via a button press when the white fixation dot changed to red. The central dot changed color to either red, green, or blue semi-randomly at a rate of approximately 30 per minute.

### MRI data acquisition

All functional and structural images were acquired using a 3 Tesla Siemens Prisma scanner at the Dartmouth Brain Imaging Center at Dartmouth College. A 32-channel receive-only phased array head coil was used for all data acquisition.

*T1 images:* For each participant, a high-resolution whole-brain anatomical volume was collected using a T1-weighted magnetization-prepared rapid acquisition gradient echo (MPRAGE) sequence (1 mm isovoxel resolution; 208 sagittal slices; TR = 2,300 ms; TE = 2.03 ms; flip angle = 9^◦^; FOV = 256 x 256 mm, bandwidth = 240 Hz/px).

*Functional images:* Two types of functional scans were acquired across two sessions, both using a T2*-weighted gradient-recalled echoplanar imaging multiband pulse sequence. Across all scans, slices were oriented approximately in plane with the calcarine sulcus to ensure coverage of most, and often all, of the temporal, parietal, and occipital lobes. At the beginning of each session, a pair of EPI images with phase encoding directions of opposite polarity in the anterior-to-posterior plane were collected for post-hoc correction of EPI spatial distortion. Session 1 consisted of localizer scans designed to identify category-selective regions of interest (nominal spatial resolution = 2 mm x 2 mm x 2mm; 69 oblique slices; TR = 2,000 ms; TE = 30 ms; flip angle = 79^◦^; matrix size 106 x 106; field of view = 212 mm x 212 mm; phase partial Fourier scheme of 6/8; bandwidth = 1,814 Hz/px; echo spacing = 0.66 ms; excite pulse duration = 8,200 microseconds; multiband acceleration factor = 3; phase encoding direction = AP). Session 2 comprised the event-related retinotopy scans and therefore used a scanning protocol that prioritized a faster TR to sample the hemodynamic response more finely (nominal spatial resolution = 2 mm x 2 mm x 2mm; 69 oblique slices; TR = 1,350 ms; TE = 33.60 ms; flip angle = 68^◦^; matrix size 106 x 106; field of view = 212 mm x 212 mm; phase partial Fourier scheme off; bandwidth = 2358 Hz/px; echo spacing = 0.54 ms; excite pulse duration = 5,140 microseconds; multiband acceleration factor = 4; phase encoding direction = AP)

### Stimulus display and scanner peripherals

Stimuli were presented using a Panasonic PT-DW750BU 7000-Lumen WXGA DLP Projector and an SR Research in-bore back-projection screen positioned just inside the head end of the magnet. Use of an in-bore screen allowed for a stimulus that subtended a greater visual angle. Participants viewed the screen via a mirror mounted atop the head coil. The viewing distance was 127 mm from the eye to the mirror + 1,010 mm from the mirror to the screen = 1,137 mm total. The maximum extent of the image projected on the screen was 297 mm x 297 mm. This resulted in a maximum possible visual angle of 14.97 ° square.

A Lenovo Thinkpad T480s computer running Linux Ubuntu 20.04.6 (Focal Fossa) controlled the stimulus presentations and recorded button presses using Matlab R2022a and extensions from PsychToolbox version 3.0.18.

To verify accurate and steady eye fixation during the retinotopic mapping runs, eye movements were recorded using an SR Research Eyelink 1000 eyetracker mounted underneath the back projection screen. Eye tracking was performed for the right eye, and samples were obtained at 250 Hz using the Pupil-CR centroid mode with default thresholding. Black gaffer tape was applied to the eye cutouts of the headcoil in an effort to reduce unwanted reflections from the infrared illuminator. Before each run, a calibration, validation, and drift correction was performed, and data was discarded if the results of the calibration were poor. During data collection, the experimenters monitored real-time plots of eye position shown on the Eyelink host computer located inside the MRI control room. Experimenters intervened and restarted data collection if eye fixation was not maintained steadily enough throughout the run. The eyetracking data, however, was low-quality for some participants. Notably, achieving sufficient pupil contrast was more difficult for participants who required MR-compatible corrective lenses. In addition, the nose bridge on the head coil sometimes partially occluded the pupil for participants with narrow inter-pupillary distance. High-quality eye-tracking data was obtained from 14/20 controls and 14/20 DPs. Based on visual inspection of the eye gaze recordings, one control participant was discarded due to insufficiently stable eye fixation.

Behavioral responses from scanning sessions were collected using a Current Designs two-button fiber optic handheld response pad connected to a Current Designs 932 interface.

### Pre-processing of MRI data

All DICOM images were converted to NIfTI format using dcm2niix version 1.0.2018.11.25 ***(Li et al., 2016)***. From each participant’s T1-weighted volume, cortical surface meshes of the pial/gray matter boundaries and gray matter/white matter boundaries were reconstructed using Freesurfer’s *recon-all* program ***(Fischl and Dale, 2000)***. Functional data from the retinotopy scans were temporally resampled with cubic interpolation to correct for differences in slice time acquisition and to simultaneously upsample the time resolution from 1,350 ms to 1,000 ms with a custom Matlab script.

Subsequent pre-processing was performed using AFNI version 23.3.12 Septimius Severus ***(Cox, 1996)***. AFNI’s *afni_proc.py* was used to create processing pipelines for both the localizer and retinotopy scans. Slice timing differences were corrected, and non-linear geometric distortion correction was applied using separately acquired volumes in which the phase-encoding direction was reversed (‘blip-up/blip-down’ strategy). EPI-to-anatomical alignment was carried out using the lpc+ZZ cost function while checking for possible left-right flips ***(Reynolds et al., 2023)***. Motion correction was applied, and timepoints were censored when the Euclidean norm (*enorm*) exceeded 0.3 mm or when the outlier fraction exceeded 5%. Spatial blurring of FWHM = 4 mm (twice the EPI voxel dimensions) was applied to the localizer data but omitted from preprocessing of the retinotopy data. Volumetric data from each scan was then projected onto the corresponding hemispheres of the high-density SUMA standard meshes (created from the Freesurfer output with AFNI function @SUMA_Make_Spec_FS). Finally, each time series was scaled to units of local BOLD percent signal change. All pre-processing results were carefully visually inspected to ensure quality control using the QC HTML reports generated by *afni_proc.py* ***(Reynolds et al., 2023)***.

### pRF analysis

The population receptive field (pRF) was modeled using the compressive spatial summation (CSS) model ***(Kay et al., 2013a)***, which was fit independently to each voxel’s preprocessed timecourse. The CSS model fits a 2D isotropic Gaussian to each voxel of the form:

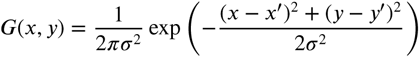

with parameters *x*^′^ and *y*^′^ describing the center coordinates of the pRF, and *σ*, the standard deviation of its Gaussian profile. The Gaussian is normalized to the unit area to make the amplitude of the response more interpretable as a percent increase.

The input stimulus to the CSS model consists solely of the spatial extent (not content) of the flickering imagery revealed through the aperture or mask at each timepoint. Therefore, the stimuli for the purpose of model fitting were prepared by converting the aperture masks to binarized (contrast) images with values including 0 (no contrast) and 1 (full contrast). Masks were downsampled to 200 by 200 pixels and then concatenated across time. The predicted response at each timepoint was obtained by computing the overlap (via dot product) between each stimulus mask and the 2D Gaussian, then multiplying it by a non-negative scaling factor, *g* for gain, and exponentiating the result by a parameter *n* constituting a static power-law nonlinearity. Afterward, the resulting time series was convolved with a model of the hemodynamic response function. The predicted BOLD response at each timepoint, *B*(*t*), can therefore be expressed formally as:

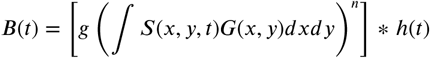

where *S*(*x*, *y*, *t*) is the contrast image at timepoint *t*, *G* is the 2D isotropic Gaussian, and ℎ(*t*) is the hemodynamic response function used in SPM. The static power-law nonlinearity, *n*, is included because it captures the nonlinear spatial summation and position tolerance of pRFs, particularly in the later stages of the visual hierarchy, in a single parameter that is easy to interpret ***(Finzi et al., 2021; Kay et al., 2013a; Le et al., 2017)***. The parameter *n* is typically observed to be less than 1 making it a *compressive* exponent.

After fitting the five pRF parameters (*x*, *y*, *σ*, *g*, *n*) in the CSS model, the Cartesian coordinates, (*x*, *y*), of the pRF centers were converted to polar coordinates to produce maps of polar angle, 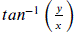, and eccentricity, 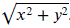. pRF size was defined as 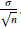. Lastly, pRF size and center location were converted from pixel units to degrees of visual angle.

Model fitting was performed using the Levenberg-Marquardt approach to nonlinear parameter estimation (MATLAB function *lsqcurvefit()* from the Optimization Toolbox). The initial seed for the solver was chosen by pre-computing model predictions for 4,632 grid points, with different combinations of *x*, *y*, *σ*, and *n*, and picking the point that produces the highest correlation with the observed data. To avoid local minima, a two-stage refined fitting procedure was implemented. In the first stage, we optimized the *x*, *y*, *σ*, and *g* parameters with the *n* parameter, controlling the compressive nonlinearity, fixed at one of three values (0.5, 0.25, or 0.125, chosen based on the prior grid search). In the second stage, the seed was updated to the solution found in the first stage, and then all five parameters were optimized simultaneously. Lower bounds of 0 and 0.1 were placed on the *σ* and *n* parameters, respectively. The maximum allowed iterations was 500, and the step tolerance was 1*e* − 6. Data from all five runs was averaged together in time prior to fitting pRFs.

### Eye tracking analysis

For participants for whom eye movements were successfully recorded (14/20 controls and 14/20 DPs), the time series of horizontal and vertical eye positions were obtained and parsed by the proprietary Eyelink software. All pre-processing steps were performed separately by run. First, time periods detected by the Eyelink software as blinks were removed along with an additional 300 msecs on either shoulder. Censored datapoints were then interpolated using a method based on discrete cosine transforms ***(Wang et al., 2012)***, and the resultant time series were low-pass zero-phase filtered using a second-order Butterworth filter with cutoff frequency of 15 Hz. Then, the data were detrended using a linear and quadratic polynomial to account for slow drifts in gaze position. Finally, the data were median-centered, the blink periods were recensored, and the time series were downsampled to 50 Hz. Units for all eye fixations were converted from pixel coordinates on the display screen to degrees of visual angle.

Fixation performance for each subject and run was summarized by computing the area of the 95% probability contour (in units of degrees of visual angle squared) encircling the distribution of eye positions.

## Data Availability

The raw fMRI recordings from both the functional localizer and retinotopy experiments are available via OpenNeuro at https://openneuro.org/datasets/ds006472.

## Code Availability

The code used to preprocess the raw fMRI timeseries and fit pRFs to the resultant timecourses is available via Figshare at https://doi.org/10.6084/m9.figshare.29582888.v1.

## Acknowledgments

We are truly grateful to all our research participants, whose dedication and enthusiasm was essential to this work. We extend our heartfelt thanks to Logan Dowdle for his generous support in developing and testing the fMRI pulse sequences used. We also deeply appreciate the thoughtful guidance and encouragement provided by Kalanit Grill-Spector, Edward Silson, and Ivan Alvarez throughout various stages of the project. This work was made possible by the generous support of the National Institutes of Health (NIH) under grants 1R01EY030613-01A1 to Bradley Duchaine and R01EY034118 to Kendrick Kay.

## Supplementary Figures and Tables

**Appendix 0—figure 5.**
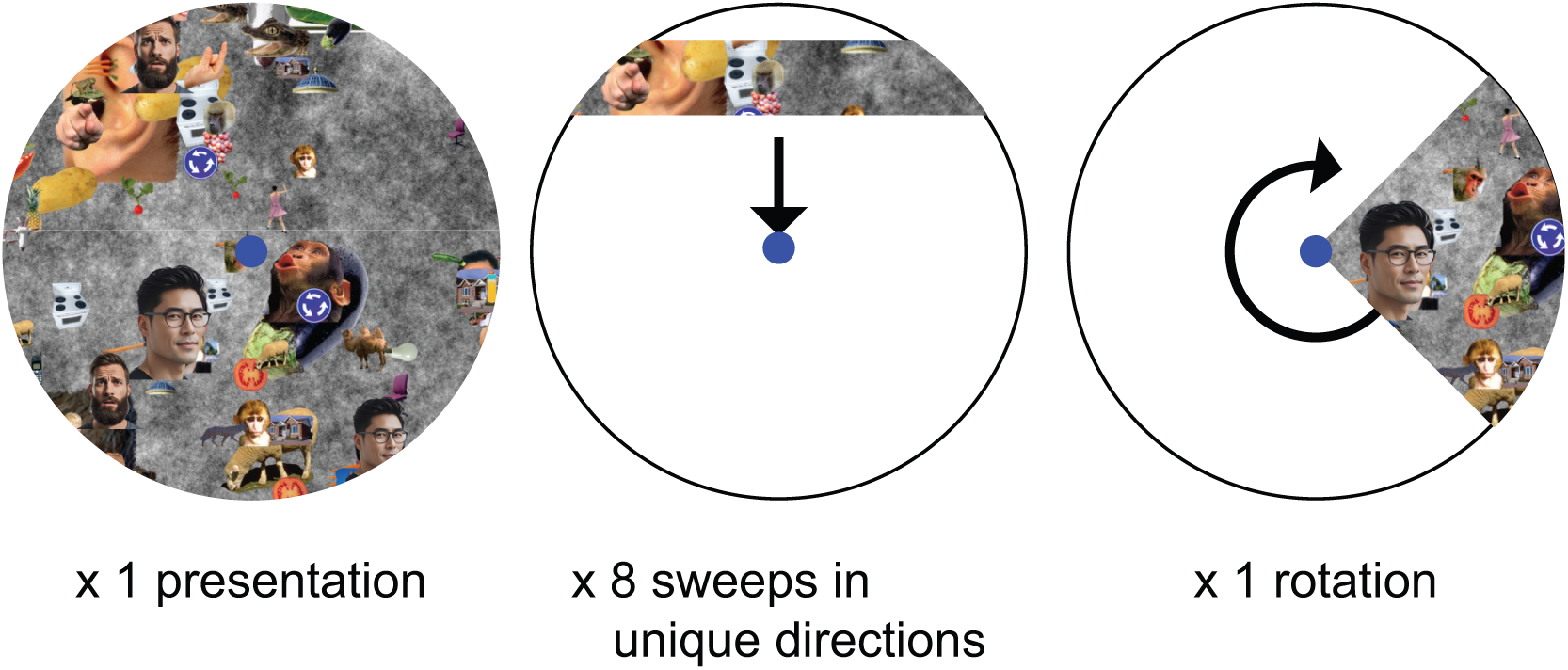
Schematic diagram of retinotopic mapping runs. Diagram is for illustration purposes only and not drawn to scale. All original faces have been replaced with AI-generated faces to preserve identifying information of original people. Runs began with a 4 second full-field exposure followed by eight sweeps of bar stimuli in eight different directions and ending with a single clockwise rotation of a wedge. Participants fixated a central dot throughout each run and indicated via button press whenever the dot changed color to red. The carrier images shown inside each aperture changed every 250 msecs (5 Hz), alternating between natural outdoor scenes and collages of various visual objects on backgrounds of pink noise (shown). See Methods for more details.

**Appendix 0—figure 6.**
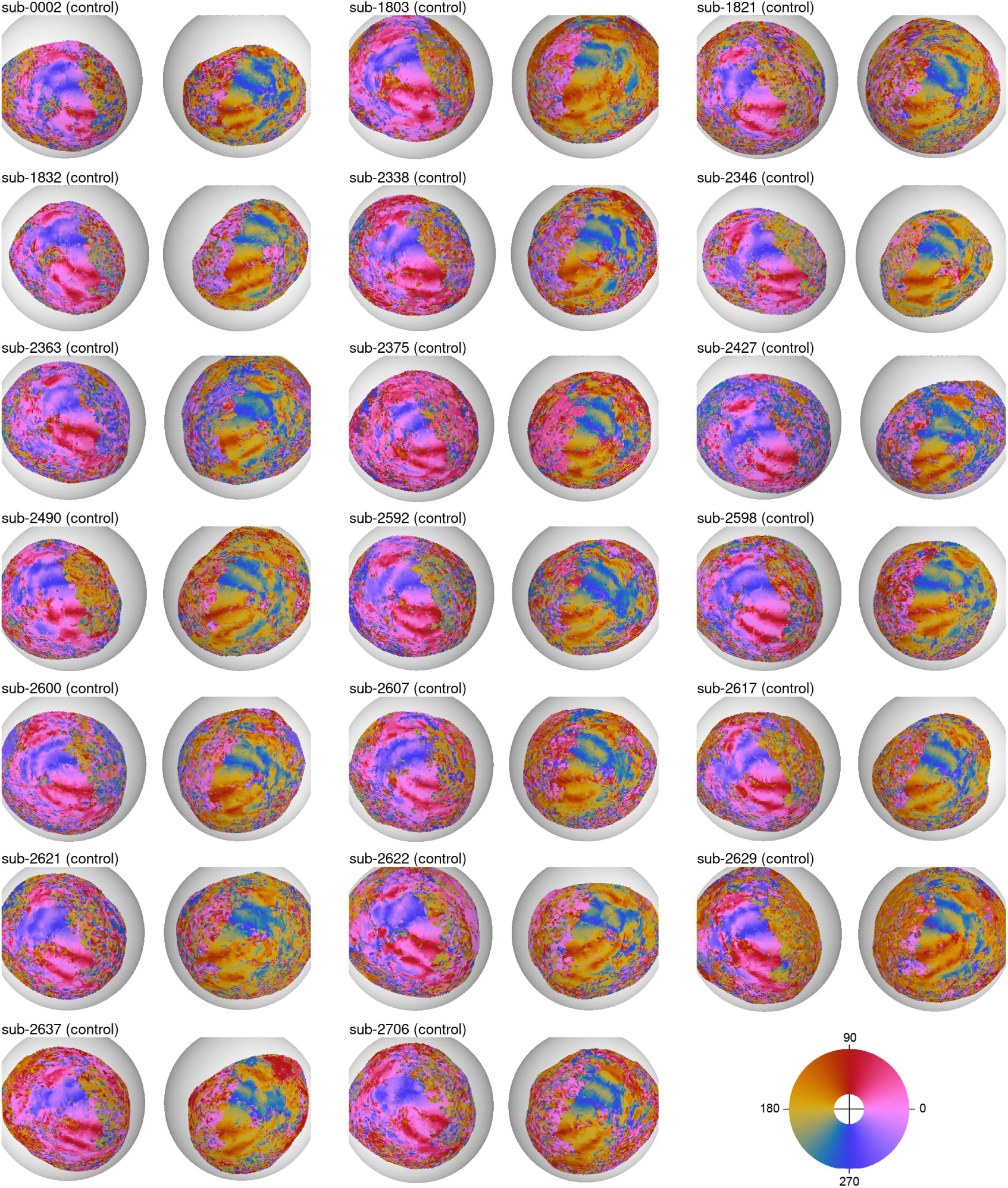
Polar angle plots displayed on inflated cortical surfaces for all control participants.

**Appendix 0—figure 7.**
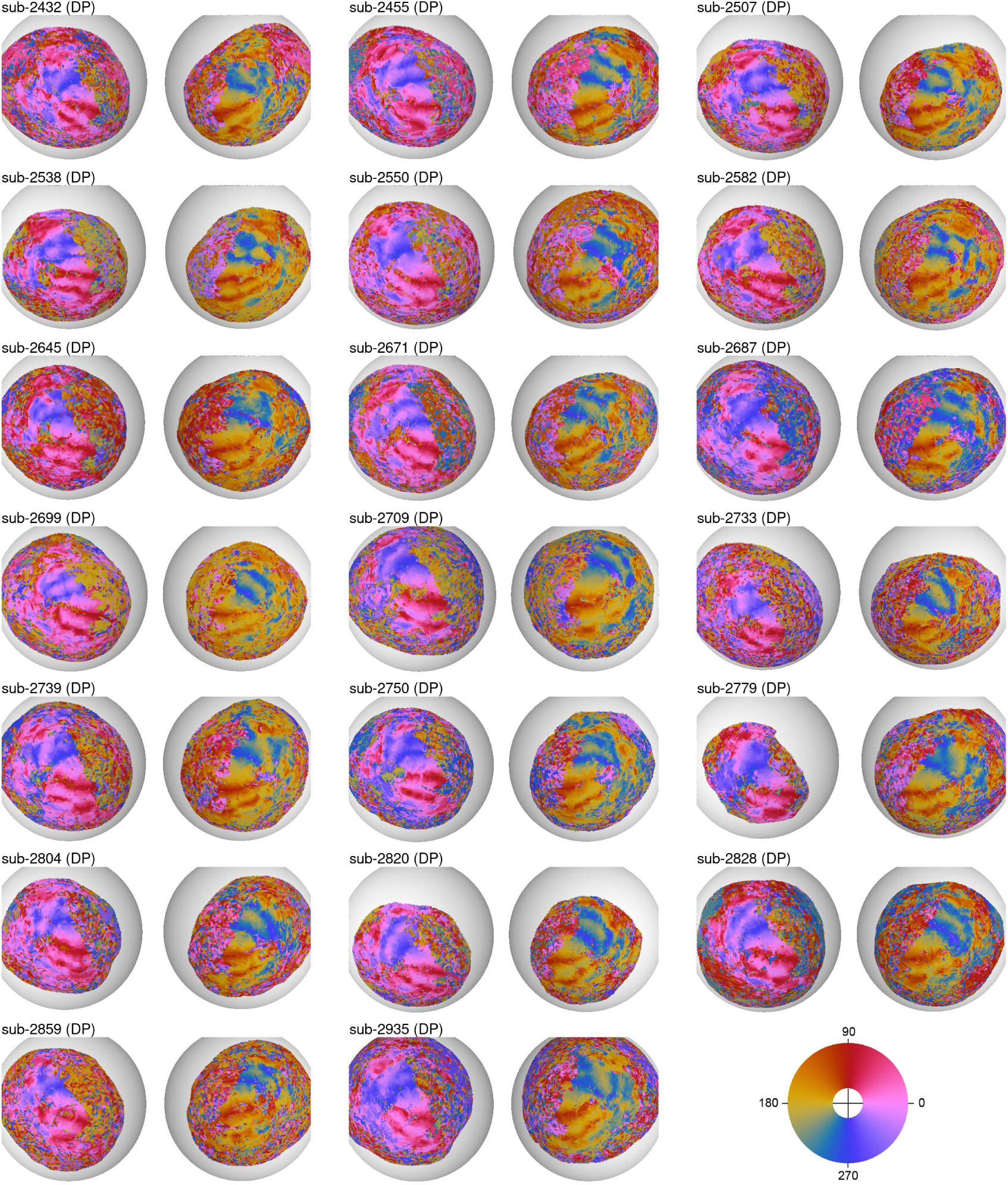
Polar angle plots displayed on inflated cortical surfaces for all DP participants.

**Appendix 0—figure 8.**
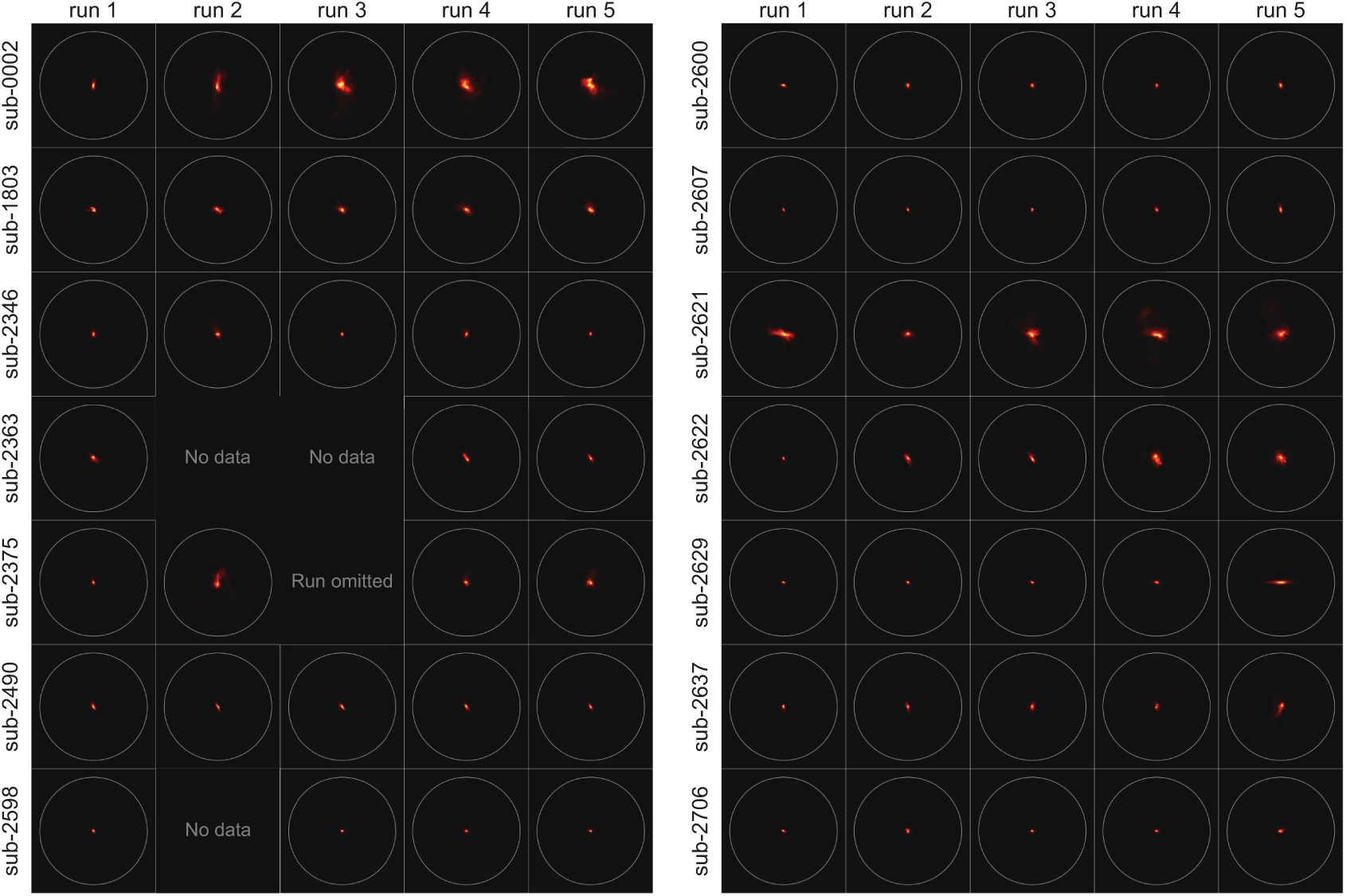
Two dimensional density plots (heatmaps) of eye position recordings by subject (controls) and run.

**Appendix 0—figure 9.**
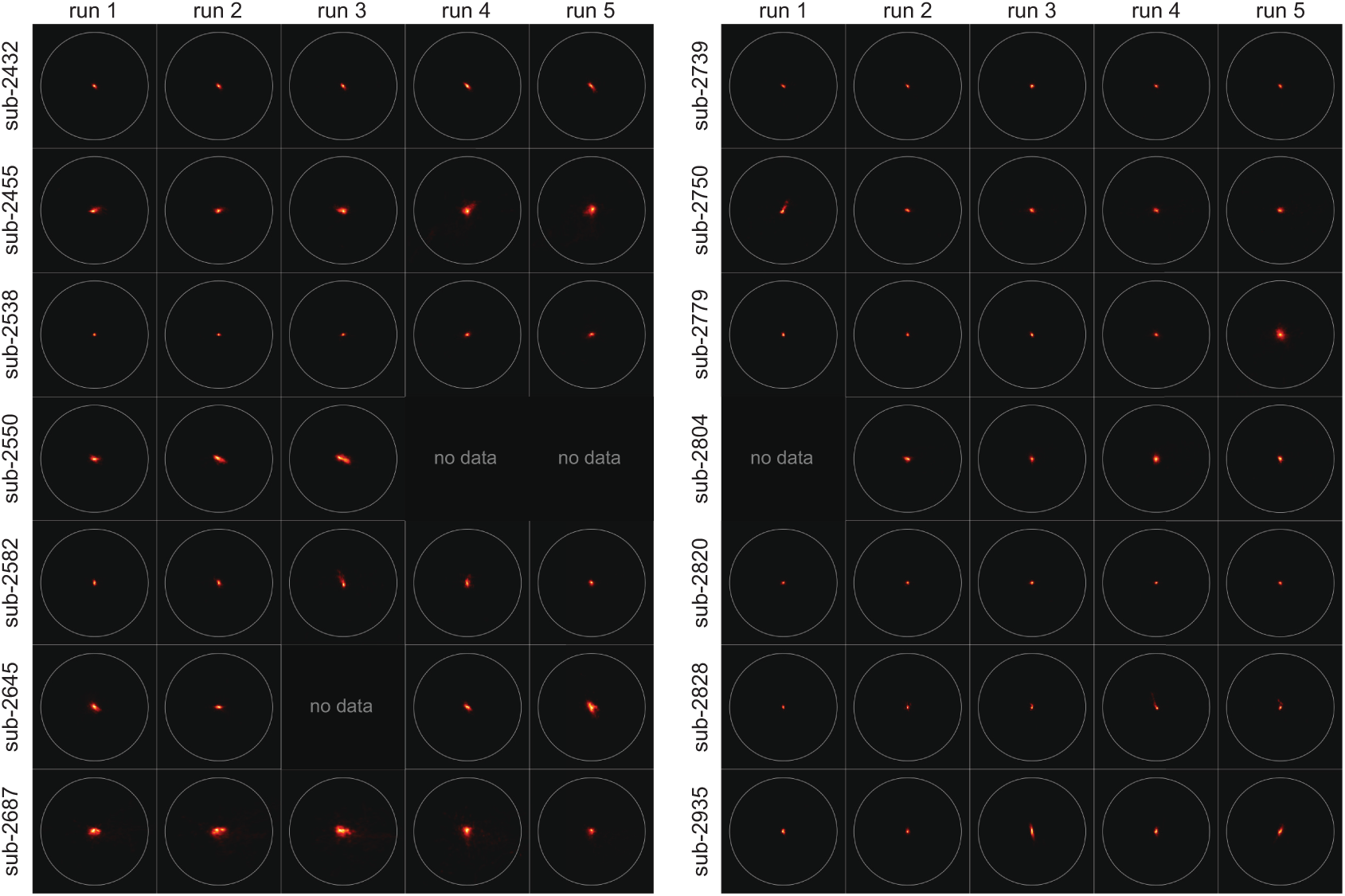
Two dimensional density plots (heatmaps) of eye position recordings by subject (DPs) and run.

**Appendix 0—figure 10.**
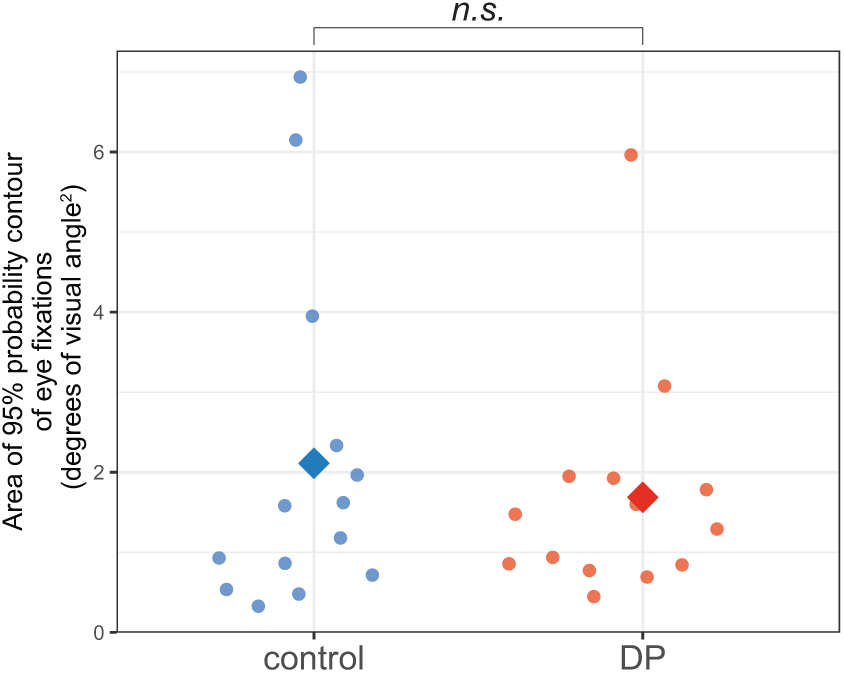
Eye fixation performance during the retinotopic mapping experiment for controls and DPs. Eye fixation performance was summarized by computing the area of the 95% probability contour of eye position samples. Points are averages across all runs obtained. Diamonds are averages for each group (Controls, DPs).

**Appendix 0—figure 11.**
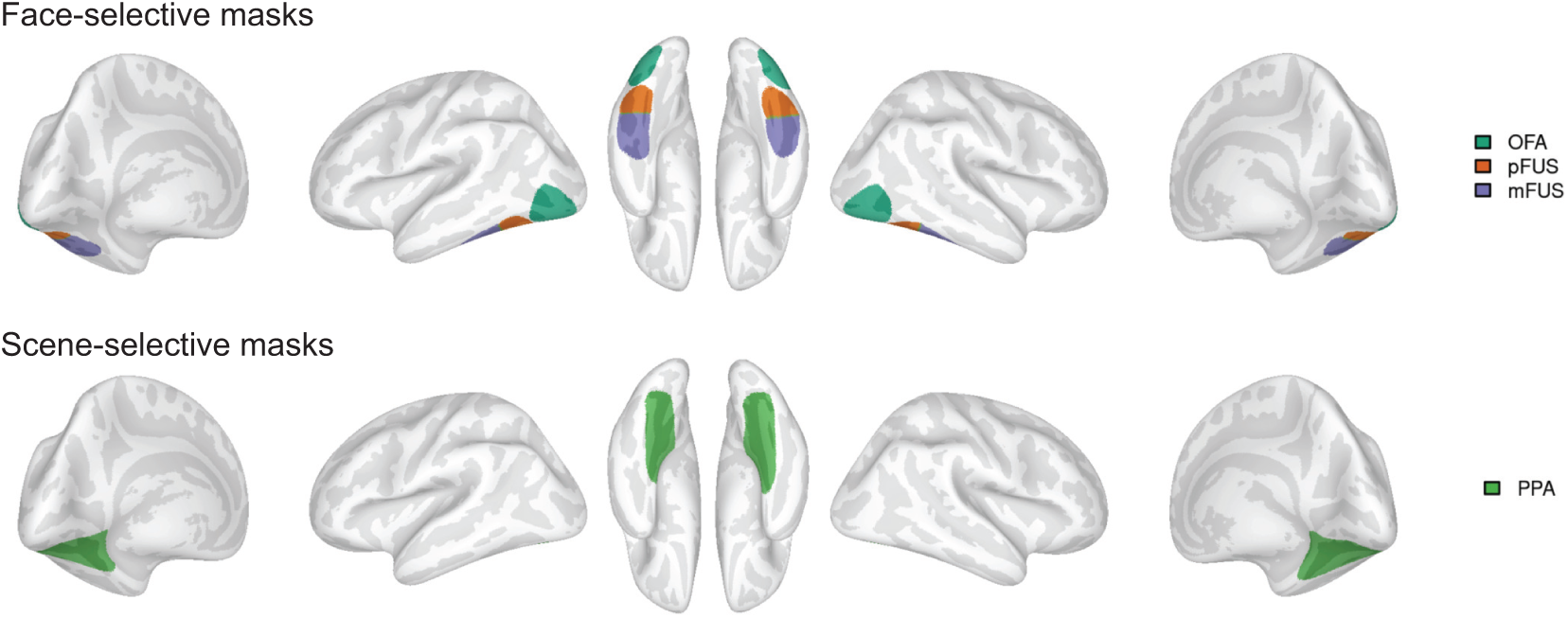
Group-derived ROI masks used for the variable window analysis.

**Appendix 0—table 3.** Category selectivity by ROI. The preferred category is defined as faces for face-selective regions (OFA, pFUS, mFUS) and scenes for scene selective regions (PPA). Category-selectivity was defined as the percent signal change to blocks of the preferred stimuli minus the percent signal change to blocks of object stimuli. To quantitatively compare category-selectivity in DPs and controls, we used the variable window method (Norman-Haignere et al., 2013) whereby voxels were selected for the analysis by applying group-defined region of interest (ROI) masks and choosing the top 20% of the voxels with the highest *t* value for the relevant contrast (Faces vs Objects for face-selective regions and Scenes vs Objects for scene-selective PPA). Voxel selection was fully cross-validated in a leave-one-run-out fashion – voxels were selected based on three out of four runs and face selectivity was measured from the left out run. For each participant, the final measure of face-selectivity was the average across four cross-validated folds. For each ROI and hemisphere, separate Welche two sample *t*-tests were conducted comparing data from DPs to controls. Due to the number of tests conducted, a more conservative alpha threshold of *p* < 0.01 was used to establish statistical significance. ∗ *p* < 0.01; ∗∗ *p* < 0.001

| ROI | hemisphere | Preferred category (PSC) |  | Objects (PSC) |  | *t* | df | *p* | Cohen's *d* |  |  |
| --- | --- | --- | --- | --- | --- | --- | --- | --- | --- | --- | --- |
|  |  | Control | DP | Control | DP |  |  |  | lower | effect | upper |
| OFA | right | 2.33±0.08 | 2.09±0.07 | 1.08±0.06 | 1.27±0.06 | 3.28 | 34.75 | 0.002* | -1.72 | -1.06 | -0.39 |
| OFA | left | 1.72±0.09 | 1.51±0.06 | 0.89±0.06 | 1.06±0.06 | 2.87 | 32.36 | 0.007* | -1.58 | -0.93 | -0.27 |
| pFUS | right | 2.65±0.08 | 2.42±0.07 | 1.24±0.08 | 1.25±0.05 | 1.56 | 35.32 | 0.1280 | -1.13 | -0.51 | 0.13 |
| pFUS | left | 2.13±0.06 | 1.81±0.07 | 1.09±0.06 | 1.19±0.05 | 3.06 | 36.09 | 0.004* | -1.64 | -0.99 | -0.33 |
| mFUS | right | 2.06±0.07 | 1.62±0.08 | 0.64±0.05 | 0.63±0.04 | 2.36 | 37.84 | 0.0240 | -1.40 | -0.77 | -0.12 |
| mFUS | left | 1.62±0.08 | 1.31±0.05 | 0.56±0.04 | 0.5±0.03 | 1.70 | 28.41 | 0.1000 | -1.18 | -0.55 | 0.08 |
| PPA | right | 2.03±0.09 | 2.19±0.07 | 1.1±0.06 | 1.04±0.05 | -1.84 | 37.53 | 0.074 | -0.04 | 0.60 | 1.23 |
| PPA | left | 1.92±0.07 | 2.07±0.06 | 0.95±0.05 | 0.98±0.05 | -1.14 | 37.89 | 0.263 | -0.26 | 0.37 | 0.99 |

**Appendix 0—table 4.** Fixed effect parameter estimates for models evaluating pRF model goodness-of-fit, *R*^2^, by hemisphere (right, left) and group (control, DP). Separate models were created for each region of interest. In each model, the intercept has been mapped to the control group in the left hemisphere. Formula (R, lme4 package): *R*^2^ ~ group + hemisphere + group:hemisphere + (1|subjectID/hemisphere)

|  | Estimate | SE | df | t | p |
| --- | --- | --- | --- | --- | --- |
| **V1** |  |  |  |  |  |
| (Intercept) | 45.50 | 1.43 | 49.27 | 31.91 | 0.000 |
| group (DP) | -2.09 | 2.02 | 49.28 | -1.04 | 0.305 |
| hemisphere (right) | -0.85 | 0.93 | 40.02 | -0.92 | 0.362 |
| group:hemisphere | 0.82 | 1.31 | 40.03 | 0.63 | 0.535 |
| **V2** |  |  |  |  |  |
| (Intercept)1 | 46.26 | 1.27 | 49.72 | 36.40 | 0.000 |
| groupDP1 | -1.31 | 1.80 | 49.72 | -0.73 | 0.470 |
| hemiRH1 | 1.46 | 0.84 | 40.08 | 1.73 | 0.092 |
| groupDP:hemiRH1 | -0.09 | 1.19 | 40.11 | -0.08 | 0.939 |
| **V3** |  |  |  |  |  |
| (Intercept) | 46.85 | 1.43 | 49.48 | 32.77 | 0.000 |
| group (DP) | -0.04 | 2.02 | 49.49 | -0.02 | 0.984 |
| hemisphere (right) | 1.70 | 0.94 | 39.86 | 1.81 | 0.078 |
| group:hemisphere | -0.42 | 1.33 | 39.89 | -0.32 | 0.754 |
| **hV4** |  |  |  |  |  |
| (Intercept) | 50.66 | 2.19 | 53.36 | 23.13 | 0.000 |
| group (DP) | -0.02 | 3.10 | 53.36 | -0.01 | 0.995 |
| hemisphere (right) | 2.54 | 1.69 | 39.85 | 1.51 | 0.140 |
| group:hemisphere | -0.74 | 2.39 | 39.84 | -0.31 | 0.757 |
| **OFA** |  |  |  |  |  |
| (Intercept) | 31.50 | 2.07 | 54.49 | 15.25 | 0.000 |
| group (DP) | 5.01 | 2.92 | 54.49 | 1.72 | 0.092 |
| hemisphere (right) | 2.14 | 1.64 | 39.97 | 1.30 | 0.201 |
| group:hemisphere | -0.95 | 2.32 | 39.97 | -0.41 | 0.686 |
| **pFUS** |  |  |  |  |  |
| (Intercept) | 27.45 | 2.17 | 71.57 | 12.63 | 0.000 |
| group (DP) | 2.49 | 3.07 | 71.53 | 0.81 | 0.421 |
| hemisphere (right) | 0.54 | 2.49 | 40.03 | 0.22 | 0.828 |
| group:hemisphere | -3.30 | 3.52 | 40.05 | -0.94 | 0.354 |
| **mFUS** |  |  |  |  |  |
| (Intercept) | 13.57 | 1.36 | 65.75 | 9.97 | 0.000 |
| group (DP) | -1.15 | 1.92 | 65.83 | -0.60 | 0.553 |
| hemisphere (right) | -1.57 | 1.41 | 39.95 | -1.12 | 0.271 |
| group:hemisphere | 0.45 | 2.00 | 40.10 | 0.23 | 0.822 |
| **PPA** |  |  |  |  |  |
| (Intercept) | 28.05 | 1.52 | 60.35 | 18.42 | 0.000 |
| group (DP) | -0.84 | 2.15 | 60.36 | -0.39 | 0.699 |
| hemisphere (right) | 1.14 | 1.41 | 39.90 | 0.81 | 0.423 |
| group:hemisphere | 0.85 | 1.99 | 39.89 | 0.43 | 0.671 |

**Appendix 0—table 5.** pRF to behavior correlations. Correlations between pRF measures (size, eccentricity, VFC area, and center dispersion) and behavioral scores on the Cambridge Face Memory Test (CFMT) and Famous Faces Test (FFT). Correlations were computed separately within each hemisphere and ROI. VFC area was computed at the 95% probability density contour. *df*: degrees of freedom; *r*: Pearson’s correlation coefficient; *p*: *p*-value uncorrected for multiple comparisons; *p_FDR_*: *p*-value corrected for multiple comparisons with FDR correction; *BF*_10_: Bayes Factor in favor of the alternative hypothesis that a true non-zero, linear relationship exists between the pair of variables.

| hemisphere | pRF measure | CFMT |  |  |  |  | FFT |  |  |  |  |
| --- | --- | --- | --- | --- | --- | --- | --- | --- | --- | --- | --- |
|  |  | df | r | p | p<sub>FDR</sub> | BF<sub>10</sub> | df | r | p | p<sub>FDR</sub> | BF<sub>10</sub> |
| hV4 |  |  |  |  |  |  |  |  |  |  |  |
| left | Size | 34 | -0.15 | 0.370 | 0.759 | 0.52 | 27 | 0.06 | 0.776 | 0.914 | 0.42 |
| left | Eccentricity | 34 | -0.24 | 0.154 | 0.588 | 0.89 | 27 | -0.24 | 0.206 | 0.588 | 0.80 |
| left | VFC area | 34 | -0.17 | 0.331 | 0.945 | 0.56 | 27 | 0.14 | 0.467 | 0.965 | 0.51 |
| left | Center dispersion | 34 | -0.08 | 0.629 | 0.818 | 0.41 | 27 | -0.07 | 0.706 | 0.818 | 0.43 |
| right | Size | 34 | 0.01 | 0.943 | 0.943 | 0.37 | 27 | 0.13 | 0.500 | 0.792 | 0.49 |
| right | Eccentricity | 34 | 0.02 | 0.891 | 0.891 | 0.37 | 27 | 0.17 | 0.384 | 0.761 | 0.56 |
| right | VFC area | 34 | 0.01 | 0.932 | 0.990 | 0.37 | 27 | 0.11 | 0.582 | 0.968 | 0.46 |
| right | Center dispersion | 34 | 0.06 | 0.707 | 0.818 | 0.39 | 27 | 0.14 | 0.470 | 0.818 | 0.51 |
| OFA |  |  |  |  |  |  |  |  |  |  |  |
| left | Size | 34 | -0.17 | 0.309 | 0.759 | 0.58 | 27 | -0.30 | 0.109 | 0.580 | 1.20 |
| left | Eccentricity | 34 | -0.12 | 0.500 | 0.770 | 0.45 | 27 | -0.25 | 0.195 | 0.588 | 0.82 |
| left | VFC area | 34 | -0.06 | 0.726 | 0.968 | 0.39 | 27 | -0.20 | 0.289 | 0.945 | 0.65 |
| left | Center dispersion | 34 | -0.25 | 0.149 | 0.387 | 0.91 | 27 | -0.38 | 0.045 | 0.373 | 2.21 |
| right | Size | 34 | -0.36 | 0.031 | 0.580 | 2.76 | 27 | -0.16 | 0.410 | 0.759 | 0.54 |
| right | Eccentricity | 34 | -0.42 | 0.011 | 0.214 | 6.24 | 27 | -0.07 | 0.709 | 0.835 | 0.43 |
| right | VFC area | 34 | -0.28 | 0.096 | 0.945 | 1.23 | 27 | -0.07 | 0.724 | 0.968 | 0.43 |
| right | Center dispersion | 34 | -0.39 | 0.020 | 0.373 | 3.87 | 27 | -0.09 | 0.631 | 0.818 | 0.45 |
| pFUS |  |  |  |  |  |  |  |  |  |  |  |
| left | Size | 34 | -0.23 | 0.174 | 0.580 | 0.83 | 27 | -0.27 | 0.149 | 0.580 | 0.98 |
| left | Eccentricity | 34 | -0.23 | 0.177 | 0.588 | 0.81 | 27 | -0.16 | 0.418 | 0.761 | 0.54 |
| left | VFC area | 34 | -0.20 | 0.237 | 0.945 | 0.68 | 27 | -0.13 | 0.499 | 0.965 | 0.49 |
| left | Center dispersion | 34 | -0.31 | 0.067 | 0.373 | 1.58 | 27 | -0.28 | 0.140 | 0.387 | 1.02 |
| right | Size | 33 | -0.28 | 0.109 | 0.580 | 1.14 | 26 | -0.06 | 0.777 | 0.914 | 0.43 |
| right | Eccentricity | 33 | -0.37 | 0.029 | 0.289 | 2.95 | 26 | -0.11 | 0.591 | 0.788 | 0.47 |
| right | VFC area | 33 | -0.28 | 0.100 | 0.945 | 1.21 | 26 | 0.00 | 0.990 | 0.990 | 0.41 |
| right | Center dispersion | 33 | -0.10 | 0.557 | 0.818 | 0.43 | 26 | -0.05 | 0.818 | 0.818 | 0.42 |
| mFUS |  |  |  |  |  |  |  |  |  |  |  |
| left | Size | 28 | -0.16 | 0.403 | 0.759 | 0.54 | 21 | -0.18 | 0.417 | 0.759 | 0.59 |
| left | Eccentricity | 28 | -0.05 | 0.787 | 0.874 | 0.41 | 21 | 0.03 | 0.880 | 0.891 | 0.45 |
| left | VFC area | 28 | -0.21 | 0.260 | 0.945 | 0.69 | 21 | -0.16 | 0.454 | 0.965 | 0.56 |
| left | Center dispersion | 28 | -0.26 | 0.167 | 0.387 | 0.90 | 21 | -0.38 | 0.075 | 0.373 | 1.62 |
| right | Size | 25 | -0.30 | 0.130 | 0.580 | 1.09 | 21 | -0.13 | 0.547 | 0.792 | 0.52 |
| right | Eccentricity | 25 | -0.30 | 0.123 | 0.588 | 1.13 | 21 | -0.20 | 0.352 | 0.761 | 0.64 |
| right | VFC area | 25 | -0.28 | 0.156 | 0.945 | 0.97 | 21 | -0.14 | 0.531 | 0.965 | 0.53 |
| right | Center dispersion | 25 | -0.31 | 0.117 | 0.387 | 1.17 | 21 | -0.06 | 0.800 | 0.818 | 0.46 |
| PPA |  |  |  |  |  |  |  |  |  |  |  |
| left | Size | 34 | 0.05 | 0.759 | 0.914 | 0.38 | 27 | 0.02 | 0.932 | 0.943 | 0.41 |
| left | Eccentricity | 34 | -0.07 | 0.705 | 0.835 | 0.39 | 27 | -0.17 | 0.378 | 0.761 | 0.56 |
| left | VFC area | 34 | 0.01 | 0.937 | 0.990 | 0.37 | 27 | 0.01 | 0.946 | 0.990 | 0.41 |
| left | Center dispersion | 34 | -0.22 | 0.193 | 0.387 | 0.77 | 27 | -0.25 | 0.193 | 0.387 | 0.83 |
| right | Size | 34 | 0.04 | 0.824 | 0.916 | 0.38 | 27 | 0.11 | 0.555 | 0.792 | 0.47 |
| right | Eccentricity | 34 | -0.13 | 0.466 | 0.770 | 0.47 | 27 | -0.11 | 0.553 | 0.788 | 0.47 |
| right | VFC area | 34 | 0.04 | 0.839 | 0.990 | 0.38 | 27 | 0.09 | 0.652 | 0.968 | 0.44 |
| right | Center dispersion | 34 | -0.11 | 0.525 | 0.818 | 0.44 | 27 | 0.05 | 0.803 | 0.818 | 0.42 |

**Appendix 0—table 6.** Bayes Factors (*BF*_01_) evaluating the evidence for the null hypothesis (no group differences between controls and DPs) in mean pRF size across ROIs and hemispheres. Values > 1 indicate evidence favoring the null hypothesis (*H*_0_), whereas values < 1 indicate evidence favoring the alternative hypothesis (*H*_1_). Bayesian independent samples *t*-tests were conducted using a Cauchy prior on the standardized effect size at two different scales: ‘medium’ (*r* ≈ 0.707) and ‘ultrawide’ (*r* ≈ 1.414) computed via the ttestBF() function in the BayesFactor R package. The ‘ultrawide’ scale tests the data against an alternative hypothesis expecting very large effect sizes, which is specifically suited for evaluating the previous claim by Witthoft et al. (2016) that pRFs in face-selective areas are approximately three times smaller in DPs compared to controls

| ROI | hemisphere | $BF_{01}$ | |
| --- | --- | --- | --- |
|  |  | medium prior | ultrawide prior |
| V1 | left | 2.69 | 4.71 |
| V1 | right | 3.16 | 5.70 |
| V2 | left | 3.00 | 5.35 |
| V2 | right | 1.57 | 2.52 |
| V3 | left | 2.07 | 3.46 |
| V3 | right | 1.39 | 2.18 |
| hV4 | left | 1.25 | 1.93 |
| hV4 | right | 2.35 | 4.02 |
| OFA | left | 1.54 | 2.46 |
| OFA | right | 0.26 | 0.32 |
| pFUS | left | 2.07 | 3.46 |
| pFUS | right | 2.49 | 4.30 |
| mFUS | left | 2.44 | 4.16 |
| mFUS | right | 2.84 | 5.00 |
| PPA | left | 2.79 | 4.92 |
| PPA | right | 2.55 | 4.43 |

## References

1. Alvarez, I., de Haas, B., Clark, C. A., Rees, G., and Samuel Schwarzkopf, D. (2015). Comparing different stimulus configurations for population receptive field mapping in human fMRI. Frontiers in Human Neuroscience, 9(FEB):1–16.

2. Arcaro, M., Mautz, T., and Livingstone, M. (2020). Anatomical correlates of face patches in macaque inferotemporal cortex. pages 1–12.

3. Avidan, G. and Behrmann, M. (2009). Functional MRI Reveals Compromised Neural Integrity of the Face Processing Network in Congenital Prosopagnosia. Current Biology, 19(13):1146–1150.

4. Avidan, G. and Behrmann, M. (2021). Spatial Integration in Normal Face Processing and Its Breakdown in Congenital Prosopagnosia. Annual Review of Vision Science, pages 1–21.

5. Avidan, G., Tanzer, M., and Behrmann, M. (2011). Impaired holistic processing in congenital prosopagnosia. Neuropsychologia, 49(9):2541–2552.

6. Avidan, G., Tanzer, M., Hadj-Bouziane, F., Liu, N., Ungerleider, L. G., and Behrmann, M. (2014). Selective Dissociation Between Core and Extended Regions of the Face Processing Network in Congenital Prosopagnosia. Cerebral Cortex, 24(6):1565–1578.

7. Bate, S. and Tree, J. J. (2017). The definition and diagnosis of developmental prosopagnosia. The Quarterly Journal of Experimental Psychology, 70(2):193–200.

8. Bates, D., Maechler, M., Bolker, B., and Walker, S. (2015). Fitting Linear Mixed-Effects Models using lme4. Journal of Statistical Software, 67(1):1–48.

9. Baudouin, J. Y. and Humphreys, G. W. (2006). Configural information in gender categorisation. Perception, 35(4):531–540.

10. Bell, L., Duchaine, B., and Susilo, T. (2023). Dissociations between face identity and face expression processing in developmental prosopagnosia. Cognition, 238(April):105469.

11. Benson, N. C., Jamison, K. W., Arcaro, M. J., Vu, A. T., Glasser, M. F., Coalson, T. S., Van Essen, D. C., Yacoub, E., Ugurbil, K., Winawer, J., and Kay, K. (2018). The Human Connectome Project 7 Tesla retinotopy dataset: Description and population receptive field analysis. Journal of Vision, 18(13):1–22.

12. Biotti, F., Wu, E., Yang, H., Jiahui, G., Duchaine, B., and Cook, R. (2017). Normal composite face effects in developmental prosopagnosia. Cortex, 95:63–76.

13. Boekel, W., Wagenmakers, E. J., Belay, L., Verhagen, J., Brown, S., and Forstmann, B. U. (2015). A purely confirmatory replication study of structural brain-behavior correlations. Cortex, 66:115–133.

14. Brewer, A. A., Liu, J., Wade, A. R., and Wandell, B. A. (2005). Visual field maps and stimulus selectivity in human ventral occipital cortex. Nature Neuroscience, 8(8):1102–1109.

15. Button, K. S., Ioannidis, J. P., Mokrysz, C., Nosek, B. A., Flint, J., Robinson, E. S., and Munafò, M. R. (2013). Power failure: Why small sample size undermines the reliability of neuroscience. Nature Reviews Neuroscience, 14(5):365–376.

16. Calder, A. J., Keane, J., Young, A. W., and Dean, M. (2000). Configural information in facial expression perception. Journal of Experimental Psychology: Human Perception and Performance, 26(2):527–551.

17. Chatterjee, G. and Nakayama, K. (2012). Normal facial age and gender perception in developmental prosopagnosia. Cognitive Neuropsychology, 29(5-6):482–502.

18. Cox, R. W. (1996). AFNI: Software for Analysis and Visualization of Functional Magnetic Resonance Neuroimages. Computets and Biomedical Research, (29):162–173.

19. Degutis, J. (2012). Face gender recognition in developmental prosopagnosia: Evidence for holistic processing and use of configural information. Visual Cognition, 20(February 2013):37–41.

20. DeGutis, J., Bahierathan, K., Barahona, K., Lee, E. M., Evans, T. C., Shin, H. M., Mishra, M., Likitlersuang, J., and Wilmer, J. B. (2023). What is the prevalence of developmental prosopagnosia? An empirical assessment of different diagnostic cutoffs. Cortex, 161:51–64.

21. Deyoe, E. A., Carman, G. J., Bandettini, P., Glickman, S., Wieser, J., Cox, R., Miller, D., and Neitz, J. (1996). Mapping striate and extrastriate visual areas in human cerebral cortex. Proceedings of the National Academy of Sciences of the United States of America, 93(6):2382–2386.

22. Dinkelacker, V., Grüter, M., Klaver, P., Grüter, T., Specht, K., Weis, S., Kennerknecht, I., Elger, C. E., and Fernandez, G. (2011). Congenital prosopagnosia: Multistage anatomical and functional deficits in face processing circuitry. Journal of Neurology, 258(5):770–782.

23. Duchaine, B. (2011). Developmental Prosopagnosia: Cognitive, Neural, and Developmental Investigations. In Rhodes, G. and Haxby, J., editors, Oxford Handbook of Face Perception, chapter 42, pages 821–838. Oxford University Press.

24. Duchaine, B. and Nakayama, K. (2005). Dissociations of face and object recognition in developmental prosopagnosia. Journal of Cognitive Neuroscience, 17(2):249–261.

25. Duchaine, B. and Nakayama, K. (2006). The Cambridge Face Memory Test: Results for neurologically intact individuals and an investigation of its validity using inverted face stimuli and prosopagnosic participants. Neuropsychologia, 44(4):576–585.

26. Dumoulin, S. O. and Wandell, B. A. (2008). Population receptive field estimates in human visual cortex. NeuroImage, 39(2):647–660.

27. Durand, K., Gallay, M., Seigneuric, A., Robichon, F., and Baudouin, J. Y. (2007). The development of facial emotion recognition: The role of configural information. Journal of Experimental Child Psychology, 97(1):14–27.

28. Engel, S. A., Glover, G. H., and Wandell, B. A. (1997). Retinotopic Organization in Human Visual Cortex and the Spatial Precision of Functional MRI. Cerebral Cortex, 7(2):181–192.

29. Epstein, R. and Kanwisher, N. (1998). A cortical representation of the local visual environment. Nature, 392:598– 601.

30. Etzel, J. A., Valchev, N., and Keysers, C. (2011). The impact of certain methodological choices on multivariate analysis of fMRI data with support vector machines. NeuroImage, 54(2):1159–1167.

31. Finzi, D., Gomez, J., Nordt, M., Rezai, A. A., Poltoratski, S., and Grill-Spector, K. (2021). Differential spatial computations in ventral and lateral face-selective regions are scaffolded by structural connections. Nature Communications, 12(1):1–14.

32. Fischl, B. and Dale, A. M. (2000). Measuring the thickness of the human cerebral cortex from magnetic resonance images. PNAS, 2000(Track II).

33. Furl, N., Garrido, L., Dolan, R. J., Driver, J., and Duchaine, B. (2011). Fusiform gyrus face selectivity relates to individual differences in facial recognition ability. Journal of Cognitive Neuroscience, 23(7):1723–1740.

34. Gerlach, C., Klargaard, S. K., Alnæs, D., Kolskår, K. K., Karstoft, J., Westlye, L. T., and Starrfelt, R. (2019). Left hemisphere abnormalities in developmental prosopagnosia when looking at faces but not words. Brain Communications, 1(1).

35. Gomez, J., Natu, V., Jeska, B., Barnett, M., and Grill-Spector, K. (2018). Development differentially sculpts receptive fields across early and high-level human visual cortex. Nature Communications, 9(1):1–12.

36. Gomez, J., Pestilli, F., Witthoft, N., Golarai, G., Liberman, A., Poltoratski, S., Yoon, J., and Grill-Spector, K. (2015). Functionally Defined White Matter Reveals Segregated Pathways in Human Ventral Temporal Cortex Associated with Category-Specific Processing. Neuron, 85(1):216–227.

37. Griffin, J. W., Bauer, R., and Scherf, K. S. (2021). Supplemental Material for A Quantitative Meta-Analysis of Face Recognition Deficits in Autism: 40 Years of Research. Psychological Bulletin, 147(3):268–292.

38. Grill-Spector, K., Weiner, K. S., Gomez, J., Stigliani, A., and Natu, V. S. (2018). The functional neuroanatomy of face perception: From brain measurements to deep neural networks. Interface Focus, 8(4).

39. Groen, I. I., Dekker, T. M., Knapen, T., and Silson, E. H. (2022). Visuospatial coding as ubiquitous scaffolding for human cognition. Trends in Cognitive Sciences, 26(1):81–96.

40. Hagler, D. J. and Sereno, M. I. (2006). Spatial maps in frontal and prefrontal cortex. NeuroImage, 29(2):567–577.

41. Harvey, B. M., Vansteensel, M. J., Ferrier, C. H., Petridou, N., Zuiderbaan, W., Aarnoutse, E. J., Bleichner, M. G., Dijkerman, H. C., van Zandvoort, M. J., Leijten, F. S., Ramsey, N. F., and Dumoulin, S. O. (2013). Frequency specific spatial interactions in human electrocorticography: V1 alpha oscillations reflect surround suppression. NeuroImage, 65:424–432.

42. Hubel, D. H. and Wiesel, T. N. (1965). Receptive Fields and Functional Architecture in Two Nonstriate Visual Areas (18 and 19) of the Cat. Journal of neurophysiology, 28(2):229–289.

43. Jednoróg, K., Marchewka, A., Altarelli, I., Monzalvo Lopez, A. K., van Ermingen-Marbach, M., Grande, M., Grabowska, A., Heim, S., and Ramus, F. (2015). How reliable are gray matter disruptions in specific reading disability across multiple countries and languages? Insights from a large-scale voxel-based morphometry study. Human Brain Mapping, 36(5):1741–1754.

44. Jiahui, G., Yang, H., and Duchaine, B. (2018). Developmental prosopagnosics have widespread selectivity reductions across category-selective visual cortex. Proceedings of the National Academy of Sciences of the United States of America, 115(28):E6418–E6427.

45. Kaiser, D., Quek, G. L., Cichy, R. M., and Peelen, M. V. (2019). Object Vision in a Structured World.

46. Kanne, S. M., Wang, J., and Christ, S. E. (2012). The Subthreshold Autism Trait Questionnaire (SATQ): Development of a brief self-report measure of subthreshold autism traits. Journal of Autism and Developmental Disorders, 42(5):769–780.

47. Kay, K. N., Weiner, K. S., Kay, K. N., and Weiner, K. S. (2015). Attention Reduces Spatial Uncertainty in Human Ventral Temporal Cortex Attention Reduces Spatial Uncertainty in Human Ventral Temporal Cortex. Current Biology, 25(5):595–600.

48. Kay, K. N., Winawer, J., Mezer, A., and Wandell, B. A. (2013a). Compressive spatial summation in human visual cortex. J Neurophysiol, 94305:481–494.

49. Kay, K. N., Winawer, J., Rokem, A., Mezer, A., and Wandell, B. A. (2013b). A Two-Stage Cascade Model of BOLD Responses in Human Visual Cortex. PLoS Computational Biology, 9(5).

50. Kay, K. N. and Yeatman, J. D. (2017). Bottom-up and top-down computations in word– and face-selective cortex. eLife, 6:1–29.

51. Kieseler, M., Dickstein, A., Krafian, A., Li, C., and Duchaine, B. (2022). HEVA-A new basic visual processing test. Journal of Vision, 22(14):4109–4109.

52. Kieseler, M. L. and Duchaine, B. (2023). Persistent prosopagnosia following COVID-19. Cortex, 162:56–64.

53. Kriegeskorte, N., Mur, M., Ruff, D. A., Kiani, R., Bodurka, J., Esteky, H., Tanaka, K., and Bandettini, P. A. (2008). Matching Categorical Object Representations in Inferior Temporal Cortex of Man and Monkey. Neuron, 60(6):1126–1141.

54. Kuznetsova, A., Brockhoff, P., and Christensen, R. (2017). LmerTest Package: Tests in Linear Mixed Effects Models. Journal of Statistical Software, 82(13):1–26.

55. Le, R., Witthoft, N., Ben-Shachar, M., and Wandell, B. (2017). The field of view available to the ventral occipito-temporal reading circuitry. Journal of Vision, 17(4):1–19.

56. Le Grand, R., Cooper, P. A., Mondloch, C. J., Lewis, T. L., Sagiv, N., de Gelder, B., and Maurer, D. (2006). What aspects of face processing are impaired in developmental prosopagnosia? Brain and Cognition, 61(2):139– 158.

57. Levine, D. N. and Calvanio, R. (1989). Prosopagnosia: A defect in visual configural processing. Brain and Cognition, 10(2):149–170.

58. Li, X., Morgan, P. S., Ashburner, J., Smith, J., and Rorden, C. (2016). The first step for neuroimaging data analysis: DICOM to NIfTI conversion. Journal of Neuroscience Methods, 264:47–56.

59. Linka, M., Broda, M. D., Alsheimer, T., de Haas, B., and Ramon, M. (2022). Characteristic fixation biases in Super-Recognizers. Journal of Vision, 22(8).

60. Lohse, M., Garrido, L., Driver, J., Dolan, R. J., Duchaine, B. C., and Furl, N. (2016). Effective connectivity from early visual cortex to posterior occipitotemporal face areas supports face selectivity and predicts developmental prosopagnosia. Journal of Neuroscience, 36(13):3821–3828.

61. Marsh, J. E., Biotti, F., Cook, R., and Gray, K. L. (2019). The discrimination of facial sex in developmental prosopagnosia. Scientific Reports, 9(1):1–8.

62. Maurer, D., Grand, R. L., and Mondloch, C. J. (2002). The many faces of configural processing. Trends in cognitive sciences, 6(6):255–260.

63. McConachie, H. R. (1976). Developmental Prosopagnosia. A Single Case Report. Cortex, 12(1):76–82.

64. McKone, E. (2009). Holistic processing for faces operates over a wide range of sizes but is strongest at identification rather than conversational distances. Vision Research, 49(2):268–283.

65. McKone, E., Hall, A., Pidcock, M., Palermo, R., Wilkinson, R. B., Rivolta, D., Yovel, G., Davis, J. M., and O’Connor, K. B. (2011). Face ethnicity and measurement reliability affect face recognition performance in developmental prosopagnosia: Evidence from the Cambridge face memory test-Australian. Cognitive Neuropsychology, 28(2):109–146.

66. Mckone, E. and Yovel, G. (2009). Why does picture-plane inversion sometimes dissociate perception of features and spacing in faces, and sometimes not? toward a new theory of holistic processing. Psychonomic Bulletin and Review, 16(5):778–797.

67. Morsi, A. Y., Chow-wing bom, H. T., Samuel, D., Goffaux, V., Dekker, T. M., and John, A. (2024). Shared spatial selectivity in early visual cortex and face-selective brain regions. bioRxiv, pages 1–37.

68. Nasiotis, K., Clavagnier, S., Baillet, S., and Pack, C. C. (2017). High-resolution retinotopic maps estimated with magnetoencephalography. NeuroImage, 145(October 2016):107–117.

69. Norman-Haignere, S., Kanwisher, N., and McDermott, J. H. (2013). Cortical pitch regions in humans respond primarily to resolved harmonics and are located in specific tonotopic regions of anterior auditory cortex. Journal of Neuroscience, 33(50):19451–19469.

70. Palermo, R., Willis, M. L., Rivolta, D., McKone, E., Wilson, C. E., and Calder, A. J. (2011). Impaired holistic coding of facial expression and facial identity in congenital prosopagnosia. Neuropsychologia, 49(5):1226–1235.

71. Pitcher, D., Walsh, V., and Duchaine, B. (2011). The role of the occipital face area in the cortical face perception network. Experimental Brain Research, 209(4):481–493.

72. Poldrack, R. A., Baker, C. I., Durnez, J., Gorgolewski, K. J., Matthews, P. M., Munafò, M. R., Nichols, T. E., Poline, J. B., Vul, E., and Yarkoni, T. (2017). Scanning the horizon: Towards transparent and reproducible neuroimaging research. Nature Reviews Neuroscience, 18(2):115–126.

73. Poltoratski, S., Kay, K., Finzi, D., and Grill-Spector, K. (2021). Holistic face recognition is an emergent phenomenon of spatial processing in face-selective regions. Nature Communications, 12(1):1–13.

74. Quené, H. and Van Den Bergh, H. (2004). On multi-level modeling of data from repeated measures designs: A tutorial. Speech Communication, 43(1-2):103–121.

75. Ramus, F., Altarelli, I., Jednoróg, K., Zhao, J., and Scotto di Covella, L. (2018). Neuroanatomy of developmental dyslexia: Pitfalls and promise. Neuroscience and Biobehavioral Reviews, 84(July 2017):434–452.

76. Reynolds, R. C., Taylor, P. A., and Glen, D. R. (2023). Quality control practices in FMRI analysis: Philosophy, methods and examples using AFNI. Frontiers in Neuroscience, 16(January):1–36.

77. Rosenthal, G., Tanzer, M., Simony, E., Hasson, U., Behrmann, M., and Avidan, G. (2017). Altered topology of neural circuits in congenital prosopagnosia. eLife, 6:1–20.

78. Rossion, B. (2009). Distinguishing the cause and consequence of face inversion: The perceptual field hypothesis. Acta Psychologica, 132(3):300–312.

79. Sereno, M. I., Dale, A. M., Reppas, J. B., Kwong, K. K., Belliveau, J. W., Brady, T. J., Rosen, B. R., and Tootell, R. B. (1995). Borders of multiple visual areas in humans revealed by functional magnetic resonance imaging. Science, 268(5212):889–893.

80. Shah, P., Gaule, A., Sowden, S., Bird, G., and Cook, R. (2015). The 20-item prosopagnosia index (PI20): A selfreport instrument for identifying developmental prosopagnosia. Royal Society Open Science, 2(6).

81. Silson, E. H., Chan, X. A. W.-y., Reynolds, R. C., Kravitz, D. J., and Baker, X. C. I. (2015). A Retinotopic Basis for the Division of High-Level Scene Processing between Lateral and Ventral Human Occipitotemporal Cortex. J.Neurosci., 35(34):11921–11935.

82. Silver, M. A. and Kastner, S. (2009). Topographic maps in human frontal and parietal cortex. Trends in Cognitive Sciences, 13(11):488–495.

83. Song, C., Samuel, D., Kanai, R., Rees, G., Song, C., Schwarzkopf, D. S., Kanai, R., and Rees, G. (2015a). Neural Population Tuning Links Visual Cortical Anatomy to Human Visual Perception. Neuron, 85(3):641–656.

84. Song, S., Garrido, L., Nagy, Z., Mohammadi, S., Steel, A., Driver, J., Dolan, R. J., Duchaine, B., and Furl, N. (2015b). Local but not long-range microstructural differences of the ventral temporal cortex in developmental prosopagnosia. Neuropsychologia, 78:195–206.

85. Steel, A., Silson, E. H., Garcia, B. D., and Robertson, C. E. (2024). A retinotopic code structures the interaction between perception and memory systems. Nature Neuroscience, 27(2):339–347.

86. Susilo, T. and Duchaine, B. (2013). Advances in developmental prosopagnosia research. Current Opinion in Neurobiology, 23(3):423–429.

87. Susilo, T., McKone, E., Dennett, H., Darke, H., Palermo, R., Hall, A., Pidcock, M., Dawel, A., Jeffery, L., Wilson, C. E., and Rhodes, G. (2010). Face recognition impairments despite normal holistic processing and face space coding: Evidence from a case of developmental prosopagnosia. Cognitive Neuropsychology, 27(8):636–664.

88. Szinte, M. and Knapen, T. (2020). Visual Organization of the Default Network. Cerebral Cortex, 30(6):3518–3527.

89. Szucs, D. and Ioannidis, J. P. (2017). Empirical assessment of published effect sizes and power in the recent cognitive neuroscience and psychology literature. PLoS Biology, 15(3):1–18.

90. Tanaka, J. W. and Farah, M. J. (1993). Parts and wholes in face recognition. The Quarterly Journal of Experimental Psychology Section A, 46(2):225–245.

91. Towler, J., Fisher, K., and Eimer, M. (2018). Holistic face perception is impaired in developmental prosopagnosia. Cortex, 108:112–126.

92. Tsantani, M., Gray, K. L., and Cook, R. (2020). Holistic processing of facial identity in developmental prosopagnosia. Cortex, 130:318–326.

93. Turner, B. O., Paul, E. J., Miller, M. B., and Barbey, A. K. (2018). Small sample sizes reduce the replicability of task-based fMRI studies. Communications Biology, 1(1).

94. van Es, D. M., van der Zwaag, W., and Knapen, T. (2019). Topographic Maps of Visual Space in the Human Cerebellum. Current Biology, 29(10):1689–1694.

95. Verfaillie, K., Huysegems, S., De Graef, P., and Van Belle, G. (2014). Impaired holistic and analytic face processing in cong prosop Evidence from eye contingent mask window. Visual Cognition, 22(3):503–521.

96. Wandell, B. A. and Winawer, J. (2015). Computational neuroimaging and population receptive fields. Trends in Cognitive Sciences, 19(6):349–357.

97. Wang, G., Garcia, D., Liu, Y., de Jeu, R., and Johannes Dolman, A. (2012). A three-dimensional gap filling method for large geophysical datasets: Application to global satellite soil moisture observations. Environmental Modelling and Software, 30:139–142.

98. Weiner, K. S. and Grill-spector, K. (2010). Sparsely-distributed organization of face and limb activations in human ventral temporal cortex. NeuroImage, 52(4):1559–1573.

99. Winawer, J., Kay, K. N., Foster, B. L., Rauschecker, A. M., Parvizi, J., and Wandell, B. A. (2013). Asynchronous broadband signals are the principal source of the bold response in human visual cortex. Current Biology, 23(13):1145–1153.

100. Winawer, J. and Witthoft, N. (2017). Identification of the ventral occipital visual field maps in the human brain. F1000Research, 6(0):1–18.

101. Witthoft, N., Poltoratski, S., Nguyen, M., Golarai, G., Liberman, A., LaRocque, K., Smith, M., and Grill-Spector, K. (2016). Reduced spatial integration in the ventral visual cortex underlies face recognition deficits in developmental prosopagnosia. bioRxiv, pages 1–26.

102. Yin, R. K. (1969). Looking at Upside-Down Faces. Journal of Experimental Psychology, 81(1):141–145.

103. Yokoyama, T., Noguchi, Y., Tachibana, R., Mukaida, S., and Kita, S. (2014). A critical role of holistic processing in face gender perception. Frontiers in Human Neuroscience, 8(JUNE):1–10.

104. Young, A. W., Hellawell, D., and Hay, D. C. (1987). Configural information in face perception. Perception, 16:747– 759.

